# Nuclear RNA Clusters Are Dynamic Structural Entities in Huntington’s Disease

**DOI:** 10.1101/2025.11.27.691003

**Authors:** Sandra Fienko, Iulia M. Nita, Ignacio Munoz-Munoz, Gillian P. Bates

## Abstract

Huntington’s disease research has focused on a toxic protein gain-of-function as the main driver of pathology. The role of mutant *HTT* mRNA *in vivo* has been only partially studied and remains largely unexplored. Recently, we discovered that fully processed human *HTT* mRNA is retained, together with the alternatively processed *HTT1a* transcript, in RNA nuclear clusters in YAC128 mouse brains. Here, we demonstrate that these clusters were already present in the prenatal stage at day 14.5, indicating early developmental effects. Moreover, these clusters were confined to neurons, implying a neuron-specific mechanism of accumulation, and were colocalised with Prpf8, a core spliceosomal protein, suggesting potential impact on nuclear homeostasis. *HTT* nuclear RNA clusters showed remarkable dynamics, rapidly dissolving when ionic interactions were disrupted or transcription and splicing were inhibited. This malleability underscores the accessibility of *HTT* mRNA to therapeutic interventions and may guide the future design of *HTT*-targeting therapies.

## Introduction

Huntingtin’s disease (HD) is a fatal neurodegenerative disorder, caused by the expanded CAG repeat in the exon 1 of huntingtin gene (*HTT*), that translates into a polyglutamine stretch in the huntingtin protein (HTT)^1^. Our previous data indicate that in the context of expanded CAG repeat, huntingtin□pre-mRNA undergoes alternative processing to generate a small transcript comprising exon 1 and 5’ intron 1 sequences (*HTT1a*)^2^. This occurs when one of two cryptic polyadenylation (polyA) signals in intron 1 is activated. The extent of *HTT*□pre-mRNA processing is determined by CAG repeat length, with a longer repeat producing greater levels of *HTT*1a^3^. Importantly, *HTT*1a has not only been detected in numerous HD mouse models^2^ but it has also been observed in patient derived fibroblast, human peripheral blood cells and post-mortem brains from HD mutation carriers^4, 5^. *HTT*1a is directly relevant to disease pathogenesis and progression as it encodes the highly pathogenic and aggregation-prone exon 1 HTT protein (HTT1a). The toxicity of the HTT1a protein isoform is underscored by the R6/2 mouse model, which expresses precisely this noxious fragment and represents the most rapidly progressing HD model^6^. Consistently, aggregates in human *post-mortem* tissue are stained with antibodies recognising N-terminal fragments of HTT^7^.

Recently, we discovered that fully processed human *HTT* mRNA is retained, together with the *HTT1a* transcript, in RNA nuclear clusters in the brains of 2-month-old YAC128 mice^8^. A subsequent study revealed that this phenomenon also occurs in BACHD-97Q-ΔN17 HD animals and could already be observed at one month of age^9^. Together, these data indicate that nuclear retention and cluster formation are specific to human huntingtin transcripts and occur early across HD mouse models, long before overt pathology. In this study we build on prior work and provide evidence that *HTT* nuclear clusters arise early in the brain development of YAC128 mice and are already present in prenatal animals.

Huntingtin protein aggregation and its downstream detrimental effects on cellular proteostasis, mitochondrial function, axonal transport, and synaptic transmission have been extensively investigated. However, a growing literature suggests a role for mutant huntingtin RNA biology in disease pathogenesis^10–13^. First, CAGs adopt secondary conformations and repeat-containing RNAs scaffold multivalent interactions that drive length-dependent phase separation and clustering *in vitro*^14^. Second, in human HD fibroblasts aberrantly expanded *HTT* mRNAs are confined to the nucleus, sequester essential splicing proteins like MBNL1 and induce splicing defects^15^. Third, comparison of two novel HD mouse models: one expressing a translatable *HTT* transgene (HD/100Q) and the other non-translatable variant (HD/100CAG) revealed that mutant *HTT* mRNA itself caused motor impairment, marked anxiety and altered striatal gene expression, although to lesser extent than in protein-expressing HD/100Q animals^13^.

In the current study, we leveraged a range of pharmacological treatments in YAC128-derived hippocampal cultures to probe the malleability of *HTT* nuclear RNA clusters. We observed that the assembly of *HTT* RNA clusters is governed by ionic interactions which play a crucial role in RNA-protein binding and RNA folding. We demonstrated that *HTT* RNA clusters can be dissolved upon application of nucleic acid intercalators underscoring their plasticity and the plausible involvement of RNA–RNA contacts in cluster stability. Moreover, we established that unperturbed transcription nucleates these structures and that blockage of canonical splicing disrupts *HTT* nuclear RNA clusters and increases the levels of *HTT1a* in the cytoplasm, possibly exacerbating HD-related pathology. *HTT* RNA clusters are extremely dynamic, which may influence their accessibility to pharmacological and therapeutic interventions. An in-depth study of huntingtin RNA metabolism as well as the nature and identity of *HTT* RNA clusters will enhance our understanding of HD pathogenesis and inform therapeutic development.

## Results

### Huntingtin nuclear RNA clusters arise early in the mouse development

Previous studies documented that human full-length huntingtin and the alternatively processed *HTT1a* mRNAs form sizable clusters in brain cell nuclei in YAC128 and BACHD-97Q-ΔN17 HD transgenic mouse models^8,9^. This phenomenon was observed at 2 and 8 months of age in YAC128 animals^8,9^ and 1,3, 6 and 8 months of age in BACHD-97Q-ΔN17 mice^9^.

To elucidate the temporal onset of this process, we set out to understand when *HTT* nuclear RNA clusters first appear. We collected brain tissue of YAC128 mice at three different developmental stages: embryonic day E14.5 and postnatal stages P1 and P15 and subjected tissues to highly sensitive and specific RNAscope experiments. To understand whether huntingtin nuclear RNA clusters stagnate or change in the late adulthood, we included brain tissue from 12-month-old mice in the analysis.

We took advantage of previously designed probes to detect either human *HTT1a* (5′ sequences of intron 1) or *FL-HTT* (exons 14–61) (Fig. 1a). We used a novel probe to identify unprocessed *HTT* pre*-*mRNA (intron 45) (Fig. 1a) as this probe uncovered higher levels of pre-mRNA present in the clusters than previously appreciated using the *HTT*inton66 probe (Supplementary Fig. 1a).

**Figure 1.**
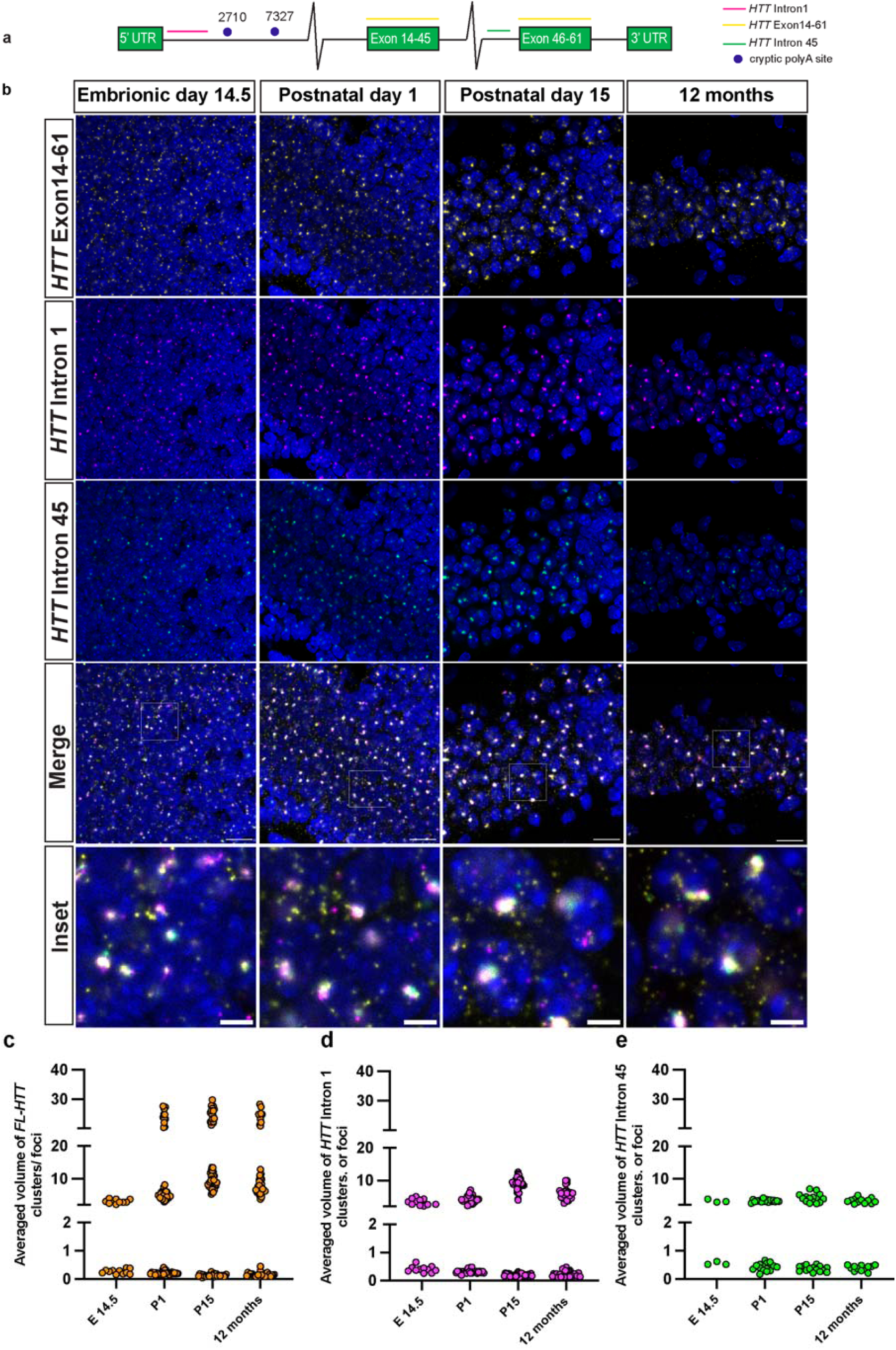
*HTT* nuclear RNA clusters arose early in the development of YAC128 mice. **(a)** Schematic showing the location of the RNAscope probes on the human *HTT* transcript: full-length *HTT* (*HTT*exon14-61), *HTT1a* (*HTT*intron 1) and pre-processed mRNA (*HTT*intron 45). **(b)** *HTT* nuclear RNA clusters composed of full-length *HTT* (yellow) and *HTT1a* (magenta) were detected already at embryonic stage 14.5 in the medial pallium, the progenitor of the hippocampus. These clusters were also observed in postnatal brains; one day or 15 days after birth as well as in the aging animals at 12 months of age. Full-length *HTT* was also detected in the extranuclear space as single transcripts. The *HTT1a* mRNA was predominantly nuclear, although single mRNA molecules were observed in the extranuclear space. The *HTT* intro*n 45* (green) probe visualised non-spliced pre-mRNA which, co-stained *HTT* nuclear clusters. Quantification of the averaged volume of **(c)** FL-*HTT* RNA, **(d)** *HTT*intron 1 or **(e)** *HTT*intron 45 clusters or transcripts at different ages of YAC128 animals. Nuclei were stained with DAPI (blue). The wild-type control sections at the corresponding ages are shown in Supplementary Fig. 1c. YAC128 (n = 3). Scale bar is 20 μm in the main image and 5 μm in the cropped magnified image.

YAC128 brain sections were hybridized with the three RNAscope probes and the medial pallium, a precursor structure that gives rise to the hippocampus, was imaged in pre- and early post-natal animals, (Supplementary Fig. 1b) or the hippocampus was imaged in the adult 12-month-old mice. The specificity of the human *HTT* probes was confirmed, as they gave no signal when hybridized to brain sections from wild-type mice (Supplementary Fig. 1c).

Using Fiji software^16^, we established a protocol to calculate volumes and numbers of nuclear clusters as well as nuclear and extranuclear single transcript molecules (Supplementary Fig. 2a). For quantification of *HTT* nuclear RNA clusters, we used an arbitrary cut-off of 2 µm^3^ to define an RNA cluster. This threshold was set based on quantification of the volume of mouse nuclear transcripts whose volume did not exceed 2 µm^3^. Single nuclear mRNA transcripts were described as puncta that ranged from 0.025 to 2 µm^3^ in volume. Single nuclear mRNAs and clusters were then binned into three separate categories based on their volume to facilitate visualisation and graphical representation as well as statistical analysis. The first bin contained large clusters between 20-50 µm^3^, the second bin contained smaller clusters between 2-20µm^3^, whereas the third bin incorporated transcripts between 0.025-2 µm^3^.

The *FL-HTT* probe revealed large RNA clusters detained in the cells of neuronal origin but not in the glial cells which mainly contained single *FL-HTT* mRNAs (Supplementary Fig. 2b). Huntingtin nuclear RNA clusters as well as single *FL-HTT* transcripts present in the nucleus and cytoplasm were detected in embryonic, postnatal and adult YAC128 mice (Fig. 1b). In early postnatal and adult mice *FL-HTT* clusters reached up to 45 µm^3^, whereas in mice at E14.5 these clusters were significantly smaller with biggest clusters approaching 17 µm^3^ in volume (Fig. 1c, Fig. 2a, Supplementary Fig. 3a). Interestingly, there was a marked increase in the volume of the smaller *FL-HTT* clusters classified in the 2^nd^ bin (2–20 µm^3^), between early postnatal age P1 and later postnatal stage P15 and then a sharp decrease in 12-month-old mice, suggesting a dynamic regulation of cluster size across developmental and aging stages (Fig. 2b). The volume of intranuclear *FL-HTT* transcripts also differed significantly between various developmental stages, with a significant decrease occurring between embryonic/P1 mice and animals transitioning to adulthood/ older mice at 12 months of age (Fig. 2c).

**Figure 2.**
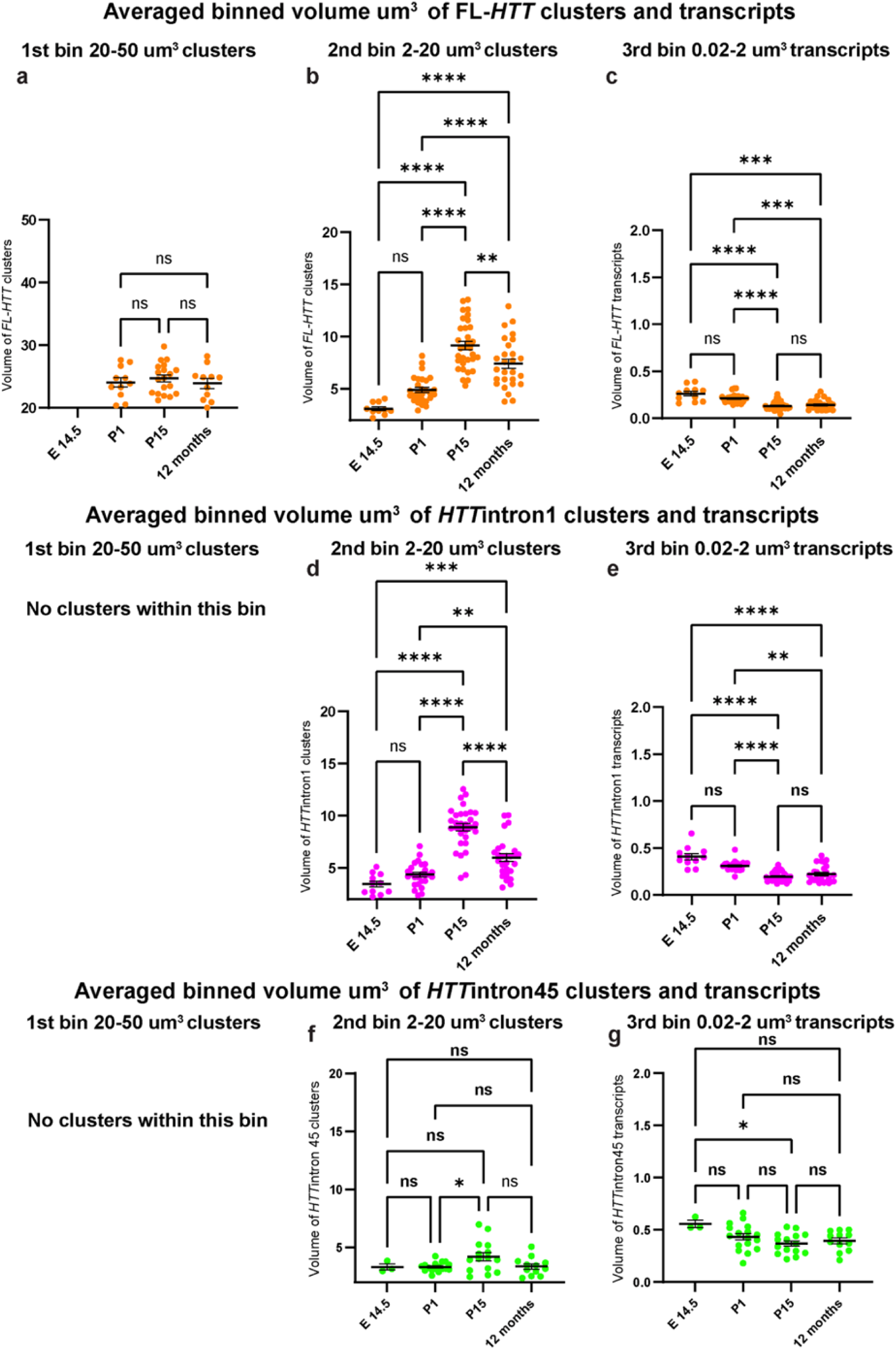
Volume of *HTT* nuclear RNA clusters was significantly changed at different developmental stages. Following quantification of the volume of *HTT* nuclear clusters and single mRNA molecules in the nucleus, results were binned into three separate categories to facilitate visual representation and statistical analysis. **(a)** Averaged volume of the FL*-HTT* clusters in the first bin remain unchanged at different developmental stages (P1=24±0.08, P15=24.7±0.05, 12-months=24±0.08). **(b)** FL*-HTT* clusters in the second bin differed significantly in volume between different ages (E14.5=3.10±0.2, P1= 4.90±0.2, P15=9.15±0.4, 12-months=7.4±0.4). **(c)** The volume of the single FL-*HTT* transcripts was also affected by development (E14.5=0.26 ±0.02, P1=0.2±0.01, P15=0.13±0.01, 12-months=0.14±0.001). **(d)** The volume of *HTT*intron 1 clusters in the second bin was impacted markedly by age (E14.5=3.5±0.3, P1= 4.4±0.2, P15=8.9±0.4, 12-months=6.0±0.4). **(e)** Changes in the volume of the single *HTT1a* molecules mirrored that of FL-*HTT* (E14.5=0.4±0.03, P1=0.3±0.01, P15=0.2±0.01, 12-months=0.2±0.02). **(f)** Averaged volume of *HTT*intron 45 clusters in the second bin differed significantly only between P1 and P15 animals (E14.5=3.3±0.3, P1=3.3±0.01, P15=4.2±0.3, 12-months=3.3±0.2). **(g)** The volume of the single *HTT*intron 45 transcripts was altered only between embryos and juvenile animals at P15 (E14.5=0.56 ±0.03, P1=0.4±0.03, P15=0.4±0.0, 12-months=0.4±0.03). FL-*HTT* (yellow), *HTT*intron 1 (magenta), *HTT* intron 45 (green), nuclei were stained with DAPI (blue). YAC128 (n = 3). Scale bar is 20 μm in the main image and 5 μm in the cropped magnified image. D’Agostino-Pearson test was applied to test data for normal distribution. The significance of data was verified either by one-way ANOVA (FL-*HTT*: 1^st^ and 2^nd^ bin, *HTTintron* 1 and *HTT*intron 45: 2^nd^ bin) or by Kruskal-Wallis test (FL-*HTT, HTTintron* 1 and *HTT*intron 45: 3^rd^ bin). Error bars = mean ± SEM, \**P* ≤ 0.033, \*\**P* ≤ 0.002, \*\*\**P* ≤ 0.0002, \*\*\*\**P* ≤ 0.0001

Strikingly, cytoplasmic *FL-HTT* mRNAs were detected exclusively as single puncta and never assembled into larger structures. Their volumes never exceeded 1 µm^3^ (Supplementary Fig. 3b) and were comparable across all ages examined, except in 12-month-old animals which showed smaller average volumes (Supplementary Fig. 3c).

As described previously, the intron 1 probe (*HTT*intron1) was present predominantly in the nucleus and co-localized with *FL*-*HTT* mRNA in the clusters^8^. *HT*Tintron1 transcripts formed clusters that did not exceeded 20 µm^3^ (Fig. 1d, Supplementary Fig. 3d). On average, the volume of the *HT*Tintron 1 clusters classified in the 2^nd^ bin was similar to the volume of the clusters formed by *FL-HTT* (Fig. 2d). Again, we observed a significant rise in the volume of *HT*Tintron 1 clusters between postnatal age P1 and later postnatal stage P15 and then a marked decrease in 12-month-old mice, which mirrored the trend showed by *FL-HTT* clusters suggesting that these clusters are regulated together. Changes in the volume of single *HT*Tintron 1 intranuclear transcripts followed those of FL-*HTT* with significant differences detected between embryonic/P1 stages and mice undergoing the transition to adulthood or older animals at 12 months (Fig. 2e).

Importantly, cytoplasmic *HTT1a* transcripts were detected in mice at all investigated stages. Quantification of the volume of extranuclear puncta revealed no significant differences between groups, except in 12-month-old animals where the decrease closely paralleled the trend associated with FL-*HTT* (Supplementary Fig.3 e,f). These results are in accordance with the *HTT1a* transcript being transported to the cytoplasm for translation to generate the HTT1a protein.

We previously reported that huntingtin nuclear RNA clusters were rarely identified by the intron 66 probe^8^ suggesting they were enriched in the spliced *HTT* mRNA. In the current study we used another intronic probe, *HT*Tintron 45 (Fig. 1a), to visualise unprocessed huntingtin mRNA. We observed nuclear clustering of *HTT*intron 45 in mice at all developmental stages (Fig. 1b,e). Clusters formed by *HTT*intron 45 were similar in volume to those formed by *FL*-*HTT* or *HTT*intron 1 transcripts for animals at E14.5 and P1 but their size remained unchanged later in adulthood (Fig .2f, Supplementary Fig. 3g). Single intranuclear pre-mRNA *HT*Tintron 45 molecules showed comparable volumes across different life stages of YAC128 animals (Fig. 2g). These data reveal a previously underappreciated presence of unprocessed pre-mRNA within the huntingtin nuclear RNA clusters.

Interestingly, volumetric analysis of mouse wild type FL*-Htt* in the nucleus revealed that these transcripts comprised a relatively homogenous population with mRNA molecules not surpassing 2 µm^3^ (Fig. 3b, c). On average, the volume of nuclear FL*-Htt* was significantly elevated in the embryos and showed no variations postnatally (Fig. 3d). Moreover, we detected a marked difference in the volume of extranuclear FL*-Htt* molecules between embryonic and juvenile mice at P15 as well as P15 and 12-month-old animals (Fig. 3e).

**Figure 3.**
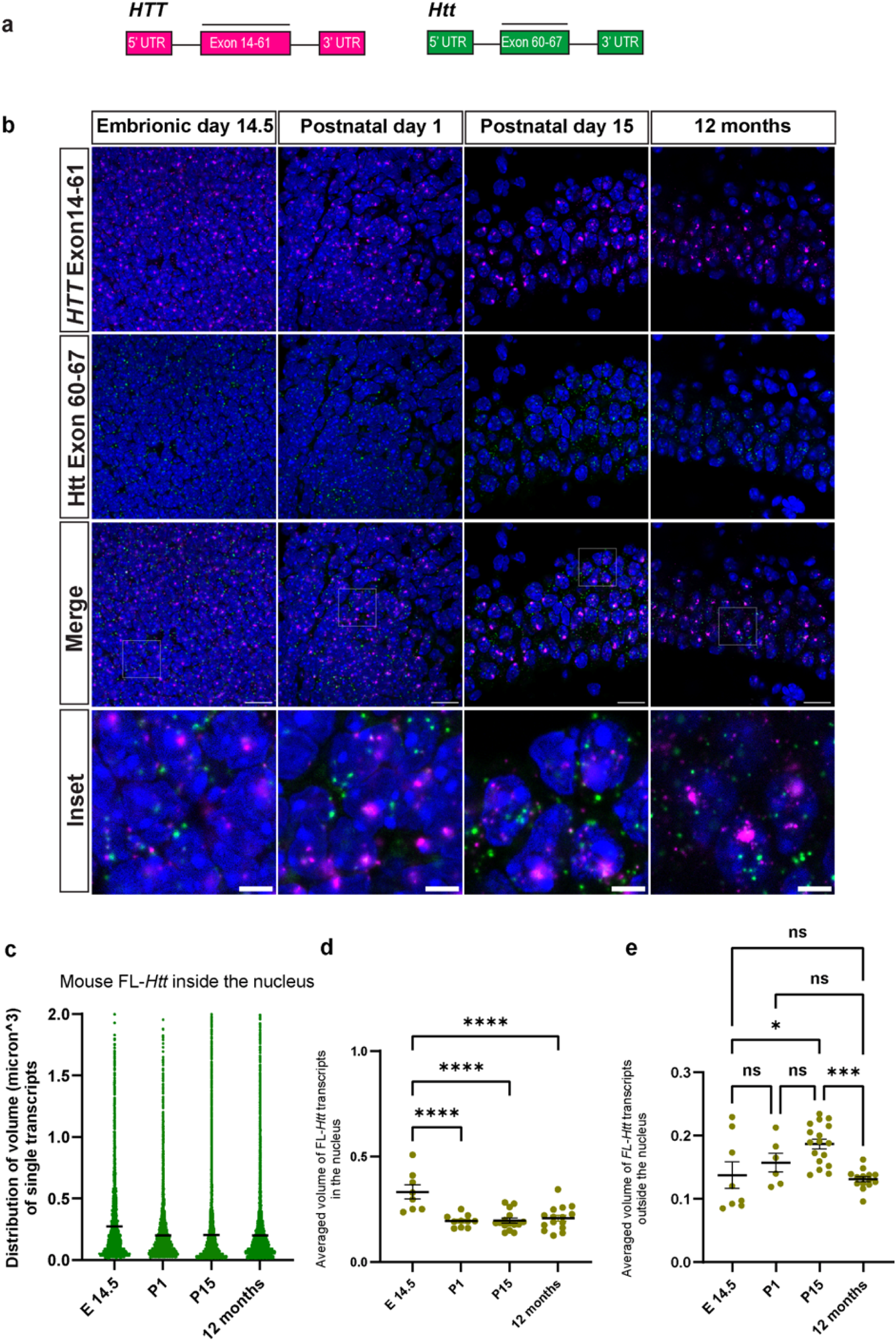
Human and mouse huntingtin mRNAs were retained in the nucleus, however only human *HTT* formed clusters. **(a)** Schematic location of the full-length human (*HTT*exon14-61) and mouse (*Htt*exon60-67) huntingtin RNAscope probes. **(b)** Full-length human *HTT* (magenta) was detected in large nuclear clusters and as single transcripts in the extranuclear space. Full-length mouse *Htt* (green) was mostly enriched in the nucleus with single molecules found in the extranuclear space. **(c)** Distribution of the volume of FL-*Htt* transcripts inside the nucleus. Quantification of the averaged volume of mouse FL-*Htt* transcripts detected **(d)** inside (E14.5=0.3±0.03, P1=0.2±0.009, P15=0.2±0.01, 12-months=0.2±0.01) or **(e)** outside of the nucleus at different ages of YAC128 mice (E14.5=0.14 ±0.02, P1=0.16±0.01, P15=0.19±0.008, 12-months=0.13±0.004). Nuclei were stained with DAPI (blue). YAC128 (n = 3). Scale bar is 20 μm in the main image and 5 μm in the cropped magnified image. D’Agostino-Pearson test was applied to test data for normal distribution and one-way ANOVA was subsequently used to verify the significance of data for the transcripts inside the nucleus, whereas Kruskal-Wallis test was applied to test the differences in the transcripts outside the nucleus. Error bars = mean ± SEM, \**P* ≤ 0.033, \*\*\**P* ≤ 0.0002, \*\*\*\**P* ≤ 0.0001

As published earlier^8^, nuclear retention and cluster formation was specific to the FL*-HTT /HTT1a* transcripts as most housekeeping mRNAs, including Ubc, *Ppib* and *Polar2a*, were primarily localized in the cytoplasm and detected at reliable levels at all ages investigated (Supplementary Fig. 4). Hybridisation with a set of bacterial probes that served as a negative control rendered no staining (Supplementary Fig. 5).

### Huntingtin RNA clusters associate with RNA splicing machinery but do not accumulate nearby well-characterized nuclear bodies

The nuclear interior is organized around chromatin into specific, microscopically visible nuclear substructures, called nuclear bodies. There are over a dozen of these organelles and each of them plays a distinct role in RNA metabolism^17^. To understand the molecular identity of *HTT* nuclear RNA clusters, we combined RNAscope and immunofluorescence (RNAscope-IF) and stained YAC128 brain sections at 2 months of age for markers of well-identified nuclear bodies. Given that these structures contain *HTT*intron 45 and are adjacent to transcription sites^9^, we hypothesised they could be localised near the splicing machinery. Evidence suggests that several proteins captured by mutant *HTT* RNA belong to the spliceosome pathway^15, 18^. Pre-mRNA-processing-splicing factor 8 (PRPF8), stood out as a pertinent candidate as it has been shown to interact with *in vitro* transcribed *HTTexon1*^18^. RNAscope-IF revealed robust colocalization of PRPF8 with *FL-HTT/HTT*intron1-rich clusters (Fig. 4a). Subsequent analysis, using the Colocalisation colormap plug-in in ImageJ software^19^, confirmed that PRPF8 interacted strongly with *HTT*intron 1 (Icor=0.65) and *FL-HTT* (Icor=0.64) nuclear clusters and that this interaction was comparable in strength to the binding between *HTT*intron 1 and FL*-HTT* (Fig. 4a, Icor=0.73).

**Figure 4.**
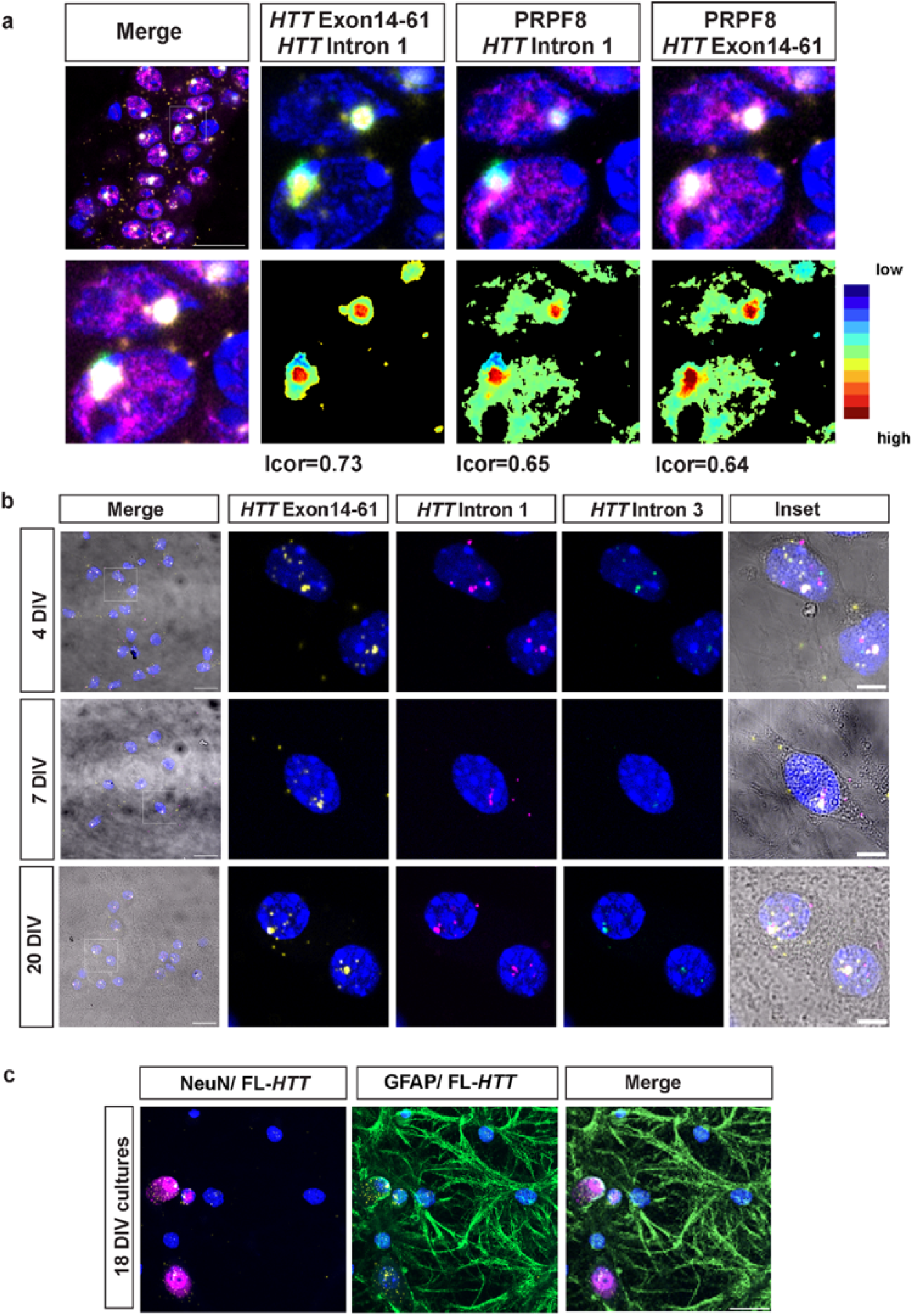
*HTT* nuclear RNA clusters colocalised with splicing machinery and were present in developing and mature hippocampal cultured neurons. **(a)** Representative images depicting that *FL-HTT* (yellow) and *HTT1a* (green) mRNAs colocalised well within the nucleus. Additionally, both transcripts colocalised with the marker of the splicing machinery (PRPF8, magenta). Icor represents the ‘Index of correlation’ and indicates the extent of colocalization. Analysis was performed using ImageJ plug-in Colocalisation Colormap. **(b)** RNAscope images showed that *HTT* nuclear RNA clusters formed as early as 4 DIV in cultured hippocampal cells and persisted when cells reached maturity (20 DIV). Full-length human *HTT*exon14-61 (yellow), *HTT*intron 1 (magenta), *HTT*intron 3 (green). Brightfield filter was added to better discriminate neuronal morphology. **(c)** Combined RNAscope-immunofluorescence experiments revealed that *HTT* nuclear RNA clusters were predominantly present in the cells of neuronal origin. GFAP, astrocytic marker (green), NeuN stained neuronal nuclei (magenta), *FL-HTT* (yellow), DAPI, nuclear stain (blue). Number of individual cultures n = 3, number of transgenic mouse pups per culture imaged n= 2. Scale bar is 20 μm in the main image and 5 μm in the cropped magnified image.

Next, we wondered whether *HTT* nuclear RNA structures were enriched within other well-identified nuclear bodies. To this end, we immunostained YAC128 sections for nuclear markers and organelles such as nucleolus (rRNA45S), Cajal bodies (coilin), paraspeckles (SFPQ), open state chromatin (H3K9ac), RNA nuclear export (p84) or different RNA-binding proteins (TDP-43, hnRNPF, hnRNPK). Analysis of the experimental data demonstrated that none of the tested markers colocalised with *HTT* nuclear RNA clusters (Supplementary Fig. 6).

These data complement previous reports that the PRPF8 splicing factor can be trapped by mutant *HTT* RNA and reiterate that this interaction might play a role in the alternative processing and production of *HTT1a*.

### Inhibitor of electrovalent bond disrupts huntingtin nuclear RNA clusters

Next, we asked whether *HTT* nuclear RNA clusters constitute malleable structures that exhibit dynamic behaviour or whether they are static. To probe this question, we took advantage of mixed primary hippocampal cultured cells derived from YAC128 newborn pups. First, we established that these cells show *HTT* nuclear RNA clusters. We performed RNAscope using three sets of probes against human *HTT1a*, FL*-HTT* or unprocessed *HTT* pre*-*mRNA (intron 3).

We interrogated developing hippocampal cells at ‘days in vitro’ (DIV) 4, cells at DIV 7 and fully matured cultures at DIV 18-20. RNAscope analysis revealed the presence of *HTT* nuclear RNA clusters already at DIV 4 that persisted at least up until DIV 20 (Fig. 4b). Moreover, RNAscope-IF confirmed that *HTT* nuclear RNA clusters were localised to neurons and not glia (Fig. 4c). Thus, mixed primary hippocampal cells constitute a reliable model to study plasticity of clusters.

Exogenously expressed repeat-containing RNA has been reported to undergo phase separation and exhibit liquid-like properties^14^. To probe whether *HTT* nuclear RNA clusters display characteristics of liquid-liquid phase separated condensates (LLPS), we took advantage of 1,6-hexanediol (Fig. 5a, Supplementary Fig. 7), and 2,5-hexanediol (Supplementary Fig. 7), agents known to hinder weak hydrophobic interactions that hold LLPS separated molecules together. Biological structures that rely upon LLPS are sensitive to 1,6-hexanediol, but less sensitive to less potent 2,5-hexanediol^17^. A 5-minute application of 8% 1,6-hexanediol to hippocampal cells did not exert any effect on the number of *HTT* nuclear RNA clusters (Fig. 5b). The number of intranuclear transcripts remained unaltered (Fig. 5c), as did the levels of cytoplasmic huntingtin mRNA molecules (Supplementary Fig. 8b). We concluded that LLPS was not a major driving force behind *HTT* nuclear RNA clusters formation. To assess whether ionic interactions affect RNA cluster stability, we applied ammonium acetate (100 mM), a cell permeable agent that does not disturb intracellular pH^14, 20^. Surprisingly, a 10-minute treatment was enough to prompt a notable decrease in the number of *HTT* nuclear RNA clusters. This decrease was significant for the FL*-HTT* and *HTT*intron 3 and showed a downward trend that did not reached significance for *HTT*intron 1 (Fig. 5d,e). Interestingly, the volume of *HTT* nuclear RNA clusters was also diminished upon NH_4_OAc treatment (Supplementary Fig. 8c), suggesting that the remaining few clusters either persist as dwarfed entities or will be dynamically dis/re-assembled later. The total number of single intranuclear mRNA huntingtin molecules was unchanged (Fig. 5f), whereas the number of cytoplasmic transcripts varied only for *HTT1a*, showing a significant drop (Supplementary Fig. 8d). These results suggest that *HTT* nuclear RNA cluster assembly is partly regulated by ionic interactions mediated by proteins that neutralize negatively charged RNA molecules.

**Figure 5.**
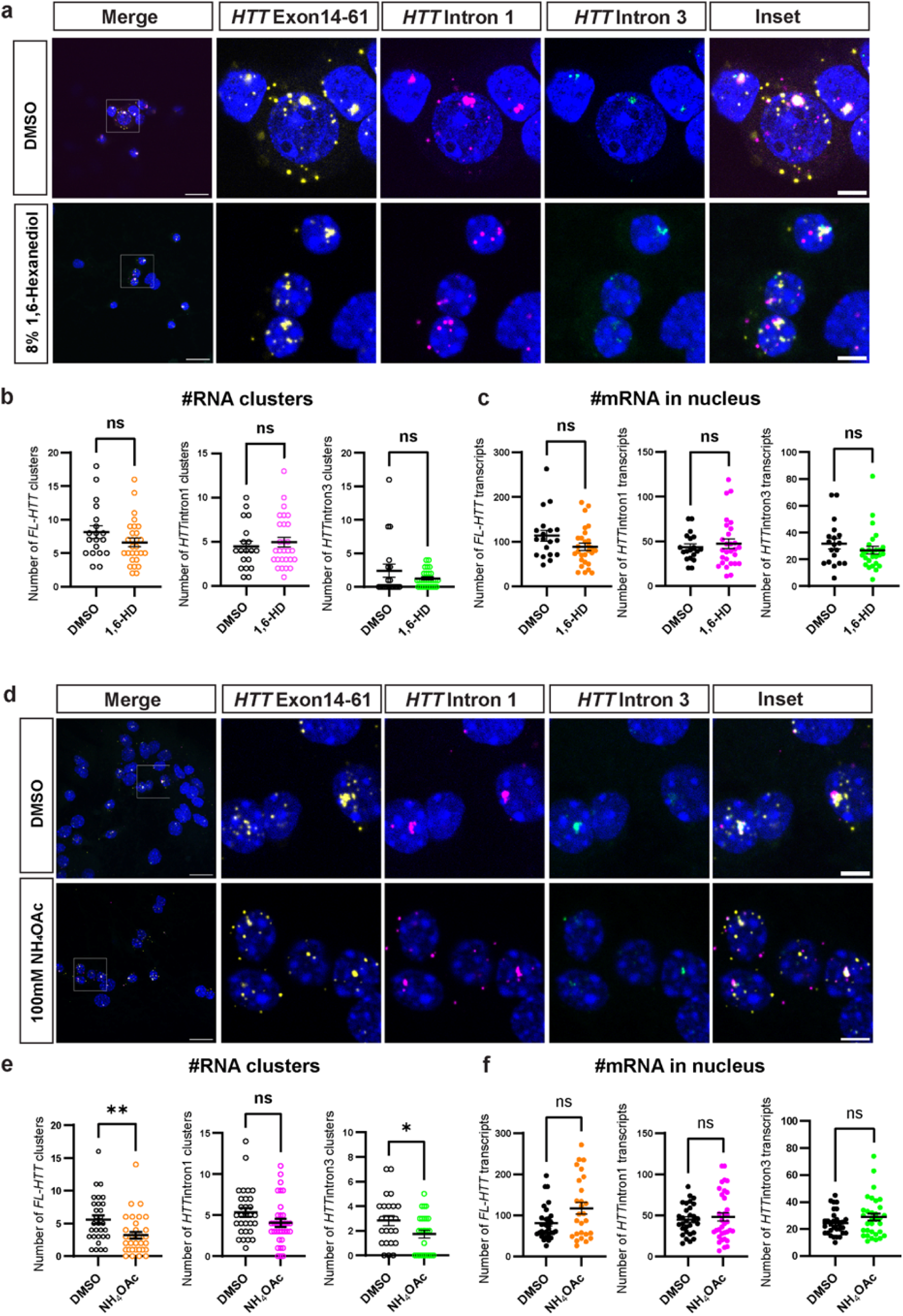
Liquid-liquid phase separation did not drive *HTT* nuclear RNA cluster formation, but these structures were disrupted by an agent that perturbs ionic interactions. **(a)** Hippocampal cultures at 18-20 DIV were treated with 8% 1,6-hexanediol or DMSO for 5 minutes. 5% 1,6-hexanediol and 8% 2,5-hexanediol were also tested and the images are shown in Supplementary Fig. 8. Quantification of **(b)** the number of *HTT* RNA clusters revealed no major changes (FL-*HTT*: 1,6-HD=6.6±0.6, DMSO=8.2±0.9; *HTT*intron 1: 1,6-HD=4.9±0.5, DMSO=4.5±0.6; *HTT*intron 3: 1,6-HD=1.2±0.2, DMSO=2.4±1. **(c)** The number of puncta in the nucleus remained unaffected (FL-*HTT*: 1,6-HD=88±8, DMSO=113±12; *HTT*intron 1: 1,6-HD=47±5, DMSO=43±3; *HTT*intron 3: 1,6-HD=27±3, DMSO=32±4). **(d)** Hippocampal cultures at 18 DIV were treated with 100 mM ammonium acetate for 10 minutes. **(e)** Quantification of the number of *HTT* RNA clusters showed significant differences in the amount of *FL-HTT* and *HTT*intron 3 nuclear clusters but not *HTT*intron 1 nuclear clusters although a trend toward lower number was observed (FL-*HTT*: NH_4_OAc=3.2±0.5, DMSO=5.6±0.6; *HTT*intron *1:*NH_4_OAc=4±0.5, DMSO=5.3±0.5; *HTT*intron 3: NH_4_OAc=1.7±0.3, DMSO=2.8±0.4). **(f)** The number of single mRNA molecules in the nucleus remain unchanged (FL-*HTT*: NH_4_OAc=117±14, DMSO=81±7; *HTT*intron *1:* NH4OAc=48±5, DMSO=45±3; *HTT*intron 3: NH_4_OAc=29±3, DMSO=24±9). FL-*HTT* (yellow), *HTT*intron 1 (magenta), *HTT*intron 3 (green), nuclei were stained with DAPI (blue). Number of individual cultures n = 3, number of transgenic mouse pups per culture imaged n= 2. Scale bar is 20 μm in the main image and 5 μm in the cropped magnified image. Shapiro-Wilk test was applied to test data for normal distribution and Mann-Whitney test was subsequently used to verify the significance of data. Error bars = mean ± SEM, \**P* ≤ 0.033, \*\**P* ≤ 0.002, \*\*\**P* ≤ 0.0002, \*\*\*\**P* ≤ 0.0001.

### Agents that intercalate into nucleic acids and block transcription induce disassembly of huntingtin nuclear RNA clusters

Next, we tested whether base-pairing interactions play a role in maintaining *HTT* RNA cluster integrity. We applied doxorubicin (2 µM), a nucleic acid intercalator, which substantially reduced the number and total volume of clusters per cell in a progressive and time-dependent manner (Fig. 6a,b; Supplementary Fig. 8e). Importantly, the number of single nuclear mRNA transcripts was already diminished for all three probes, 5 hours post-treatment and continued to decrease further (Fig. 6c). Cytoplasmic *FL-HTT* transcripts increased 5 hours after doxorubicin application to then decrease significantly 10 hours after drug treatment, *HTT1a* showed a downward time-dependent tendency (Supplementary Fig. 8f). In summary, these data suggest that intermolecular base-paring interactions contribute to stability of the *HTT* nuclear RNA clusters either by promoting intact transcription process accompanied by efficient topoisomerase activity or by preventing the efflux of a subclass of specific RBPs^21^.

**Figure 6.**
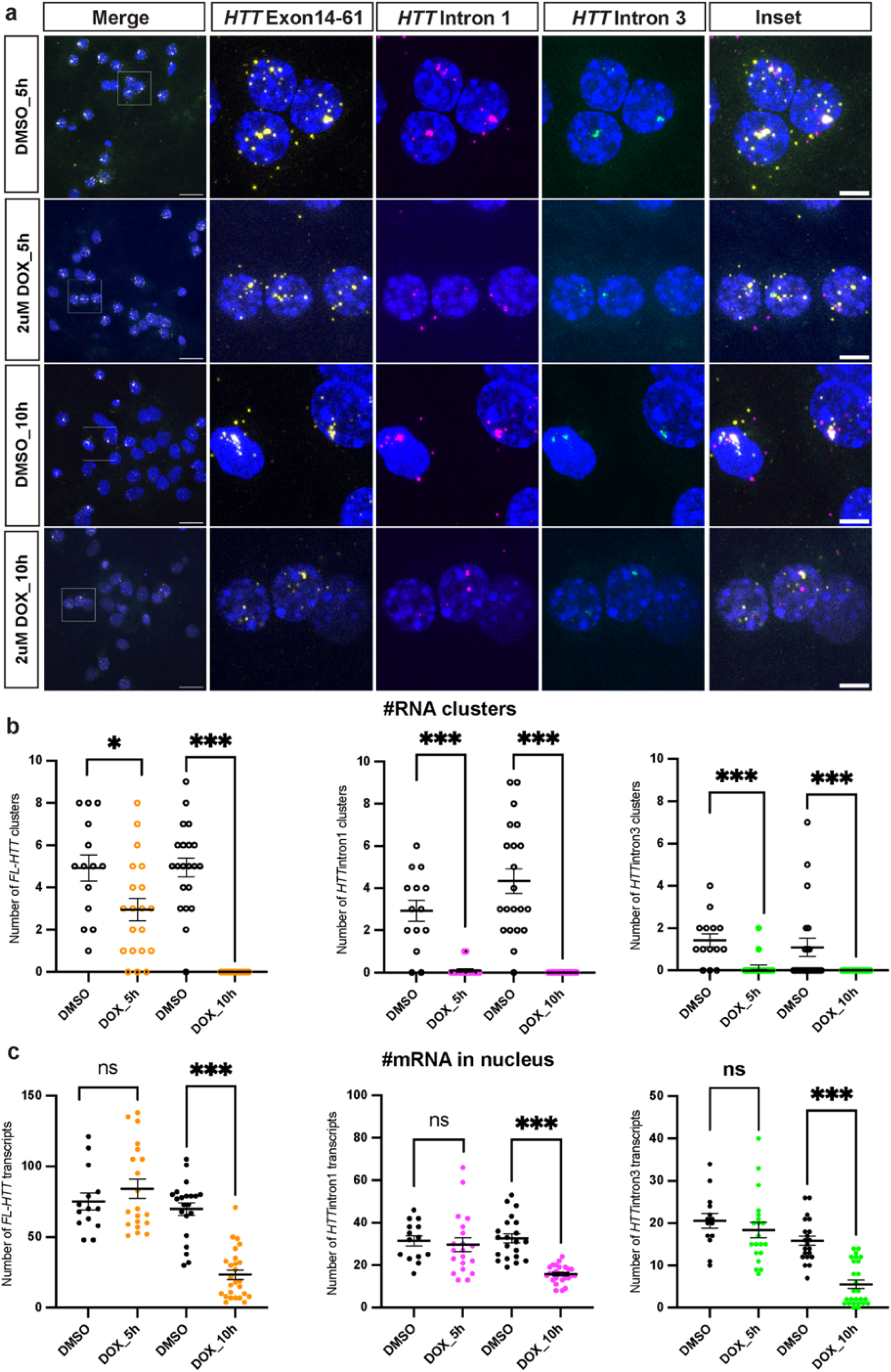
*HTT* nuclear RNA clusters disintegrate in response to doxorubicin. **(a)** 2 µM doxorubicin was applied to hippocampal cultures at 18-20 DIV either for 5 or 10 hours. **(b)** Doxorubicin diminished the number of *HTT* nuclear RNA clusters after 5 hours and completely abrogated their numbers after the 10-hour incubation (5 hours: FL-*HTT*: Dox=3±0.5, DMSO=5±0.6; *HTT*intron 1: Dox=0.1±0.07, DMSO=3±0.5, *HTT*intron 3 Dox=0.15±0.1 DMSO=1.4±0.3; 10 hours: FL-*HTT*: Dox=0.0±0.0, DMSO=5±0.45; *HTT*intron 1: Dox=0.0±0.0, DMSO=4.3±0.6, *HTT*intron 3 Dox=0.0±0.0 DMSO=1±0.4). **(c)** The number of single mRNA molecules inside the nucleus showed an initial trend toward an increase after 5 hours for FL-*HTT*, followed by significant downregulation after 10 hours (5 hours: FL-*HTT*: Dox=84±6, DMSO=75±6; 10 hours FL-*HTT*: Dox=24±3, DMSO=70±4). The levels of intranuclear *HTT* intron 1 and *HTT* intron 3 transcripts decreased progressively (5 hours: *HTT*intron 1: Dox=29.6±3 DMSO=31±2, *HTT*intron 3: Dox=18±2 DMSO=20±2; 10 hours: *HTT*intron 1:Dox=16±0.7 DMSO=33±2, *HTT*intron 3: Dox=6±1 DMSO=16±1). FL-*HTT* (yellow), *HTT*intron 1 (magenta), *HTT*intron 3 (green), nuclei were stained with DAPI (blue). Number of individual cultures n = 3, number of transgenic mouse pups per culture imaged n= 2. Scale bar is 20 μm in the main image and 5 μm in the cropped magnified image. Shapiro-Wilk test was applied to test data for normal distribution and either Mann-Whitney (5-hour treatment: number of clusters for *HTT*intron 1 and *HTT*intron 3; number of nuclear transcripts for FL-*HTT* and *HTT*intron 1; 10-hour treatment: number of clusters for all three probes; number of nuclear transcripts for FL-*HTT,* number of nuclear transcripts for *HTT*intron 3) or unpaired two-tailed *t*-test (5-hour treatment: number of clusters for FL-*HTT* and number of nuclear transcripts for *HTT*intron 3; 10-hour treatment: number of nuclear transcripts for *HTT*intron 1) was subsequently used to verify the significance of the data. Error bars = mean ± SEM, \**P* ≤ 0.033, \*\**P* ≤ 0.002, \*\*\**P* ≤ 0.0002, \*\*\*\**P* ≤ 0.0001.

Given that *HTT* nuclear RNA clusters localise near transcription sites (as revealed by colocalization of *FL-HTT/HTT1a* clusters with *HT*Tintron 45), we explored the hypothesis that intact transcription might have an impact on the integrity of *HTT* nuclear RNA clusters. We first applied the commonly used pan-transcriptional inhibitor Actinomycin D (ActD, 1 µM), which alleviated not only the number of *HTT* nuclear RNA clusters (Fig. 7a, b) but also their volume (Supplementary Fig. 8g). In parallel, ActD raised the number of mRNA molecules inside the nucleus (Fig.7c) as well as the levels of cytoplasmic mRNA puncta (Supplementary Fig.8h).

**Figure 7.**
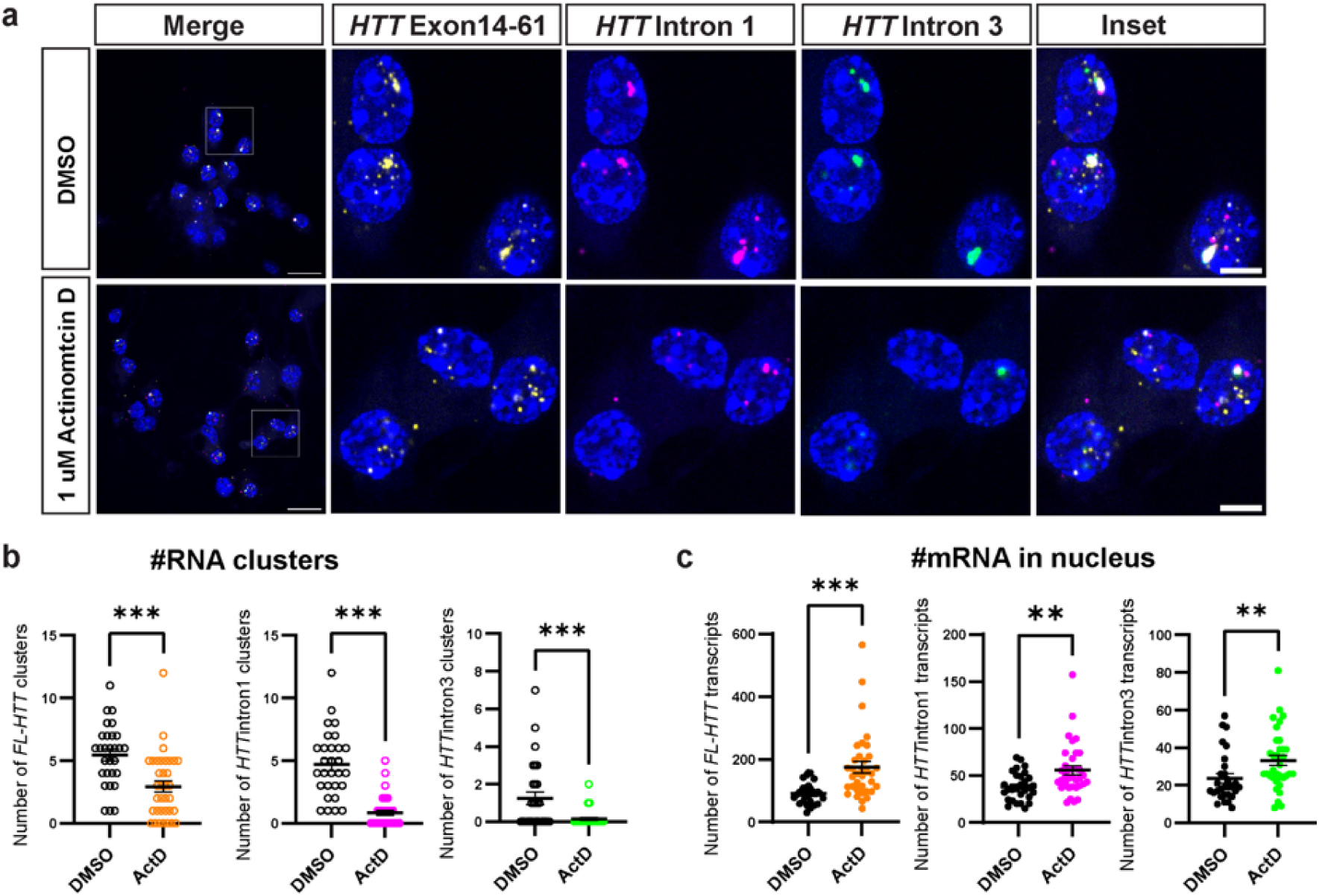
*HTT* nuclear RNA clusters were sensitive to transcription inhibition. **(a)** Hippocampal cultures at 18-20 DIV were treated with the commonly used transcription blocker 1 µM Actinomycin D (ActD) for 3 hours. **(b)** The inhibitor prevented huntingtin clustering and significantly downregulated the number of *HTT* nuclear RNA clusters (FL-*HTT*: ActD=3±0.4, DMSO=5.5±0.45; *HTT*intron 1: ActD=0.9±0.2, DMSO=4.7±0.5, *HTT*intron 3 ActD=0.14±0.07 DMSO=1.2±0.3). **(c)** The number of single mRNA molecules inside the nucleus was also notably affected (FL-*HTT*: ActD=175±2, DMSO=91±6, *HTT*intron 1: ActD=56±5 DMSO=38±3, *HTT*intron 3: ActD=33±3 DMSO=24±2). FL-*HTT* (yellow), *HTT*intron 1 (magenta), *HTT*intron 3 (green), nuclei were stained with DAPI (blue). Number of individual cultures n = 3, number of transgenic mouse pups per culture imaged n= 2. Scale bar is 20 μm in the main image and 5 μm in the cropped magnified image. Shapiro-Wilk test was applied to test data for normal distribution and Mann-Whitney test was subsequently used to verify the significance of data. Error bars = mean ± SEM, \**P* ≤ 0.033, \*\**P* ≤ 0.002, \*\*\**P* ≤ 0.0002, \*\*\*\**P* ≤ 0.0001.

### Inhibition of transcriptional elongation and pre-mRNA splicing promote dissolution of huntingtin nuclear RNA clusters

To corroborate that *HTT* nuclear RNA clusters are sensitive to inhibition of mRNA transcription and avoid off-target effects of ActD, that include DNA-intercalation, we treated hippocampal cells either with 5,6-Dichloro-1-β-D-ribofuranosylbenzimidazole (DRB,100 uM) or with NVP-2 (250 nM). Both drugs inhibit transcription elongation and block CDK9-dependent RNA Pol II activation with DRB having an additional negative effect on casein kinase II^22, 23^. Like ActD, NVP-2 and DRB caused a significant drop in the number of *HTT* nuclear RNA clusters (Fig. 8a,b,d,e).

**Figure 8.**
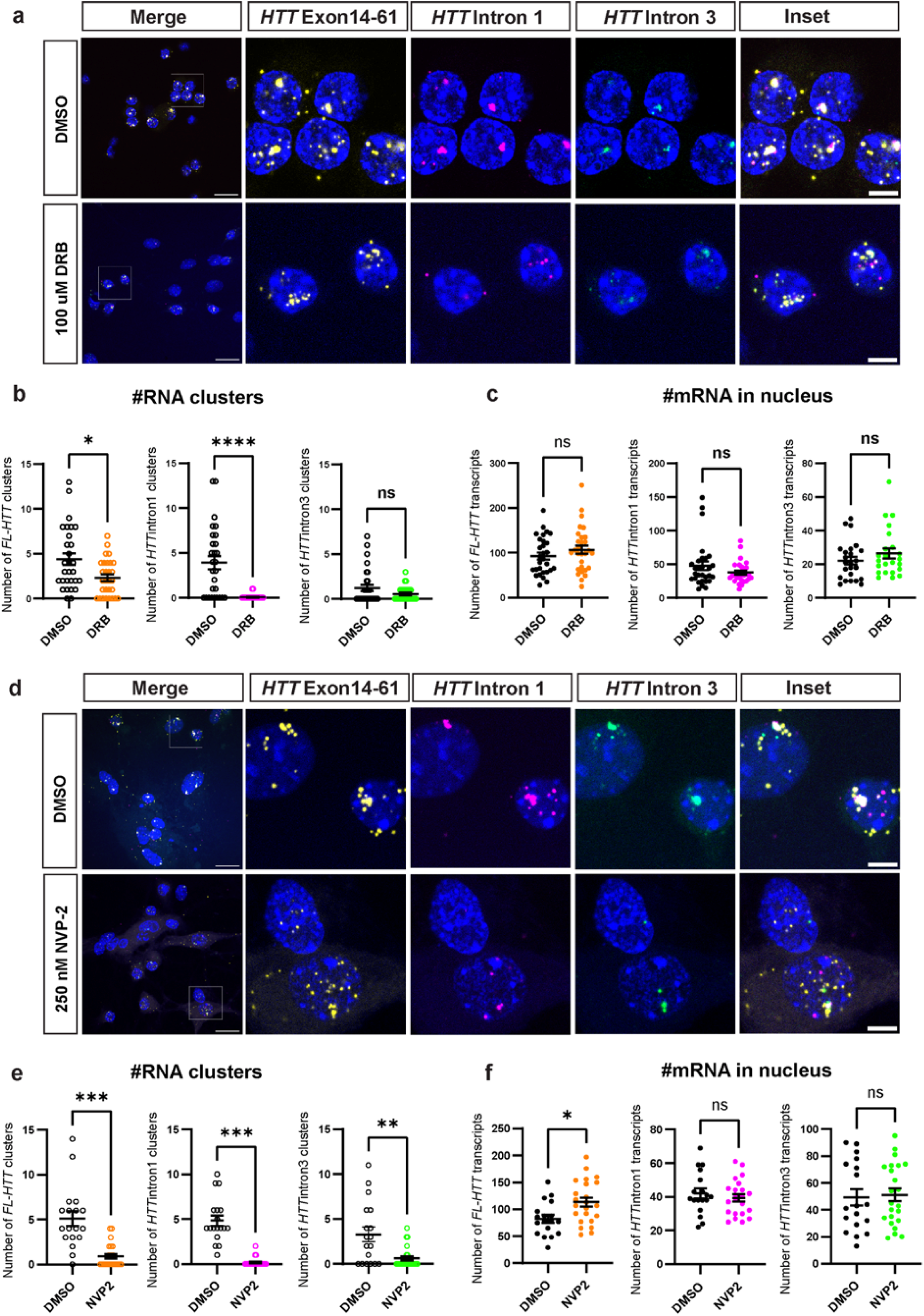
Inhibition of mRNA synthesis led to the disruption of *HTT* nuclear RNA clusters. Hippocampal cultures at 18-20 DIV were treated with transcription elongation inhibitors: **(a)** 250 nM NVP2 or **(d)** 100 µM DRB for 3 hours. **(b, e)** Quantification analysis revealed that these pharmacological agents successfully dissolved *HTT* nuclear RNA clusters (DRB:FL-*HTT*:DRB=2±0.4,DMSO=4±0.6;*HTT*intron 1:DRB=0.07±0.05,DMSO=4±0.7;*HTT*intron 3: DRB=0.5±0.1, DMSO=1.2±0.3; NVP-2: FL-*HTT*: NVP-2=1±0.1 DMSO=1.7±0.07, *HTT*intron 1: NVP-2=0.4±0.1, DMSO=1.7±0.07; *HTT*intron 3: NVP-2=1.4±0.09, DMSO=1.6±0.07). **(c)** NVP-2 had a significant effect on the number of *FL-HTT* molecules inside the nucleus without affecting numbers of either *HTT*inton 1 or *HTT*inton 3 transcripts (NVP-2: FL-*HTT*: NVP-2=113±8, DMSO=82±7, *HTT*intron 1: NVP-2=39±2, DMSO=42±3; *HTT*intron 3: NVP-2=51±8, DMSO=49±6). **(f)** The number of *FL-HTT*, *HTT*inton 1 or *HTT*intron 3 transcripts in the nucleus remained unchanged upon DBR treatment (FL-*HTT*: DRB=107±9, DMSO=93±8; *HTT*intron 1: DRB=38±3, DMSO=48.6, *HTT*intron 3: DRB=26±3, DMSO=22±2). FL-*HTT* (yellow), *HTT*intron 1 (magenta), *HTT*intron 3 (green), nuclei were stained with DAPI (blue). Number of individual cultures n = 3, number of transgenic mouse pups per culture imaged n= 2. Scale bar is 20 μm in the main image and 5 μm in the cropped magnified image. Shapiro-Wilk test was applied to test data for normal distribution and either Mann-Whitney (DRB and NVP-2 treatments: number of clusters for all three probes) or unpaired two-tailed *t*-test (NVP-2 treatment: number of transcripts in the nucleus for all three probes; DRB: number of transcripts in the nucleus for FL-*HTT*) was subsequently used to verify the significance of data. Error bars = mean ± SEM, \**P* ≤ 0.033, \*\**P* ≤ 0.002, \*\*\**P* ≤ 0.0002, \*\*\*\**P* ≤ 0.0001.

Strikingly, the total volume of FL-*HTT* nuclear RNA clusters was notably affected by both inhibitors, whereas *HTT*intron 1 clusters were severely impacted only by NVP-2, an effect that might be attributed to the greater potency of NVP-2 (Supplementary Fig. 9a,c). Interestingly, only NVP-2 produced a substantial increase in the levels of intranuclear *FL-HTT* mRNAs, without affecting the number of *HTT*intron 1 or *HTT*intron 3 mRNAs (Fig. 8c,f).

None of the drugs significantly impacted the number of cytoplasmic transcripts (Supplementary Fig .9b,d). This set of data underscores the importance of effective transcriptional elongation in the production of *HTT* nuclear RNA clusters. Our data, together with those of others^18^, implicate interaction of mutant *HTT* with components of the splicing machinery. Thus, we sought to understand whether pre-mRNA splicing plays a role in cluster formation. We used Pladienolide B (PladB, 100 nM), that targets the SF3B1 subunit in the U2 snRNP complex and inhibits early stages of the splicing reaction^23^. In concert with its effect on canonical splicing, the treatment had a significant impact on the number (Fig. 9a,b) and volume of FL-*HTT* clusters (Supplementary Fig. 9e). PladB reduced the number of *HTT*intron 1 clusters, although the effect was markedly weaker compared to that observed for FL-*HTT* (Fig. 9a,b).

**Figure 9.**
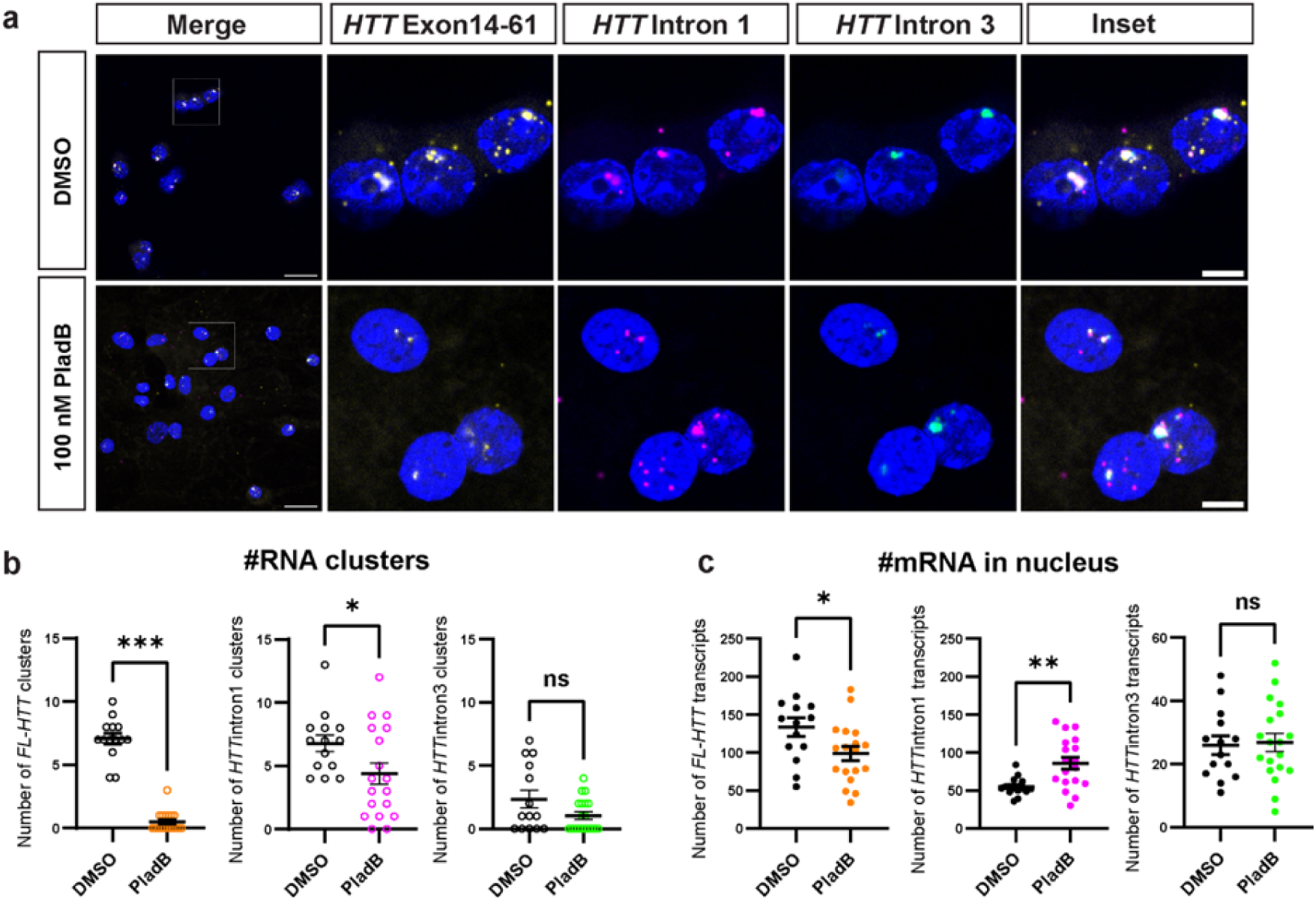
Inhibition of pre-mRNA splicing blocked formation of *HTT* nuclear RNA clusters and promoted accumulation of *HTT* intron1 mRNAs in the nucleus. **(a)** 100nM of PladB was applied to 18-20 DIV hippocampal cells for 4h. **(b)** The splicing inhibitor prompted a significant decrease in the number of *HTT* nuclear RNA clusters FL-*HTT*: PladB=0.5±0.2, DMSO=7±0.4; *HTT*intron 1: PladB=4.4±0.8, DMSO=6.8±0.6, *HTT*intron 3: PladB=1.0±0.3, DMSO=2.3±0.7), **(c)** PladB notably diminished the number of canonically spliced *FL-HTT* and significantly increased the amount of alternatively processed *HTT1a* (FL-*HTT*: PladB=99±9, DMSO=133±12, *HTT*intron 1: PladB=86±8, DMSO=55±3, *HTT*intron 3: PladB=27±3, DMSO=26±3). FL-*HTT* (yellow), *HTT*intron 1 (magenta), *HTT*intron 3 (green), nuclei were stained with DAPI (blue). Number of individual cultures n = 3, number of transgenic mouse pups per culture imaged n= 2. Scale bar is 20 μm in the main image and 5 μm in the cropped magnified image. Shapiro-Wilk test was applied to test data for normal distribution and either Mann-Whitney (number of clusters for *HTT*intron 1 and *HTT*intron 3) or unpaired two-tailed *t*-test (number of clusters for FL-*HTT,* number of transcripts in the nucleus for all three probes) was subsequently used to verify the significance of data. Error bars = mean ± SEM, \**P* ≤ 0.033, \*\**P* ≤ 0.002, \*\*\**P* ≤ 0.0002, \*\*\*\**P* ≤ 0.0001.

As inhibition of pre-mRNA splicing leads to the nuclear accumulation of either unspliced pre-mRNA or aberrantly processed mRNA and a downregulation of fully processed mRNA, we observed a significant drop in the number of single canonically spliced *FL-HTT* molecules and a substantial increase in the number of alternatively processed *HTT*intron 1 transcripts, however no effect was detected for *HTT*intron 3 (Fig. 9c). Cytoplasmic FL-*HTT* mRNA puncta were downregulated, although this trend did not reach significance, conversely the levels of *HTT1a* transcripts were markedly increased (Supplementary Fig. 9f). Together these findings imply a shift in *HTT* pre-mRNA splicing toward aberrant processing under splicing-inhibitory conditions. Moreover, elevated levels of *HTT1a* in the extranuclear space are indicative of its escape from nuclear retention and its accumulation in the cytoplasmic compartment.

## Discussion

### Mechanisms governing *HTT* nuclear RNA clusters size in developing brain

Here, we demonstrated that *HTT* nuclear RNA clusters, previously detected in YAC128 and BACHD-97Q-ΔN17 mice at 1 month of age^8, 9^, emerge prenatally as early as embryonic day 14.5. Interestingly, *FL-HTT* appeared much larger in postnatal vs embryonic animals, which was accentuated by the absence of *FL-HTT* clusters exceeding 20 um^3^ in E14.5 animals (Fig. 2a). As development progressed, the volume of the *FL-HTT* clusters rose significantly between postnatal age P1 and juvenile P15 animals before declining in 12-month-old mice.

These results might reflect differences in the transcription rate or RNA turnover; they might also stem from a divergent RNA-binding protein repertoire or differential RNA modifications that govern RNA retention in embryo vs postnatal mice. How these various processes are regulated during neuronal development remains to be elucidated. However, recent studies implicate RNA folding and RNA structural dynamics in dictating the accessibility of binding sites, spatial organization of transcripts and the recruitment of specific RBPs, all of which could impact the ability of *FL-HTT* mRNAs to assemble into nuclear clusters. The transcriptome of human embryonic stem cells (hESC) is structurally more accessible and homogeneous than that of differentiated neurons^24, 25^. This greater structural flexibility could reduce the propensity of transcripts to self-associate, resulting in smaller clusters in embryonic mice. Indeed, in response to alternations in the surrounding environment (e.g. concentration of ions, metabolites, pH, osmolarity) RNA assumes various structural states (ensembles)^26, 27^. Some of these states are thermodynamically favourable and can therefore become stabilised. These structural ensembles can interact in cis and/or trans with other RNA molecules either specifically or promiscuously, different RNA conformers can also scaffold various proteins^26^. In this way, RNA ensembles can interact with each other to drive formation of bigger *HTT* nuclear RNA clusters. By contrast, *HTT*intron 1 clusters were smaller, which is emphasized by the absence of these clusters in the first bin, containing mRNAs in 40-20 um^3^ range. This is perhaps unsurprising, as *HTT1a* is markedly smaller molecule and may adopt more compact conformation compared to FL*-HTT*. In the second bin, however *HTT*intron 1 clusters mirrored the trajectory of FL-*HTT* clusters (Fig. 2d). These similarities may reflect shared regulation by transcriptional output, splicing efficiency, RNA structural dynamics, post-transcriptional modifications, or RBP availability, all of which could impact nuclear RNA cluster assembly and stability.

Interestingly, *HTT*intron 1 is subjected to m^6^A methylation and reducing methylation levels leads to decrease in alternatively processed *HTT*1a^28^. Additionally, m^6^A modification has functional impact on many facets of RNA metabolism, including splicing, polyadenylation, nuclear export and degradation^29^. Its levels are extremely dynamic, remaining low throughout embryogenesis and rising dramatically by adulthood. The increase in the volume of *HTT*intron 1 clusters from embryonic to juvenile mice and then a drop in the older animals could reflect changes in the levels of m^6^A modification that govern splicing and nuclear export.

### *HTT* nuclear RNA clusters and splicing machinery

Our results revealed that *HTT* nuclear RNA clusters in the brains of YAC128 mice at two months of age were enriched in PRPF8. Currently, it is unclear whether these clusters exacerbate RNA-mediated toxicity or if they are protective by detaining noxious transcripts in the nucleus and thus preventing their translation.

Interaction of huntingtin mRNA with the protein that sits at the core of the splicing machinery might indicate ongoing splicing activity and suggest a regulatory role of this protein in huntingtin transcript processing. Interestingly, a genetic screen conducted in Caenorhabditis elegans identified two PRPF8 alleles that have a profound effect on cryptic splicing frequency by stabilising spliceosome’s catalytic core^30^. As the mechanism and/or participating factor(s) that promote production of *HTT1*a have not yet been elucidated, it is tempting to speculate that PRPF8 could play a central role in this process. Formation of *HTT* nuclear RNA clusters may also have deleterious effects on nuclear homeostasis by sequestering PRPF8 and presumably other factors. Indeed, PRPF8 has already been linked to polyglutamine-mediated toxicity^31^ and in the recent report, it emerged as one of 36 splicing machinery proteins that can bind to *in vitro* transcribed *HTT*exon1 RNA in a CAG repeat length-dependent manner.

### *HTT* nuclear RNA clusters are stabilised by ionic bonding

Liquid–liquid phase separation is a phenomenon whereby multivalent interactions among molecules drive their spontaneous condensation into distinct liquid phases with differing component concentrations. This process has been gaining tremendous interest as a governing force behind many cellular substructures and membraneless condensates (e.g. paraspeckles)^32^. A multitude of studies have implicated aberrant phase separation in the aetiology of neurodegeneration. It has been proposed that LLPS contributes to the interaction of disease-associated proteins which leads to the formation of highly dynamic condensates. Over time, however, these condensates can transition from dynamic to irreversible aggregates^32^. Although similar findings in HD remain limited^33^, it is appreciated that RNA with pathologically expanded repeat can also template multivalent intermolecular interactions. These interactions could lead to the formation of large clusters via LLPS. Indeed, Jain and Vale showed that exogenously expressed 47xCAG or 120xCAG RNA assemble into liquid like foci both *in vitro* and in the cell nucleus^14^. In contrast, our data revealed that *HTT* nuclear RNA clusters are not a result of phase separation. We probed LLPS by applying 1,6-hexanediol or 2,5-hexanediol, regents that disrupt phase separated condensates by impeding weak hydrophobic interactions^34^. Although, widely used, it is not fully understood how 1,6-hexanediol dissolves these interactions and it is now evident that LLPS involves other forces like electrostatic, hydrophobic, pi–pi, and pi–cation interactions^35^. While formation of *HTT* nuclear RNA clusters might not be driven by LLPS, examining varying concentrations and time durations of 1,6-hexanediol as well as testing different agents that interfere with phase separation will provide more definitive evidence.

RNA can fold into multiple secondary conformations such as hairpins, internal loops, bulges, multi-branch loops and pseudoknots^11^. Moreover, mRNAs with pathologically expanded CAG repeats form a heterogenous population of molecules with a mixture of secondary structures that are especially unstable and prone to self-associate or nucleate base-pairing with another RNA molecule^36^. Additionally, RNA can fold into tertiary structures which are exceptionally sensitive to the concentrations and types of cations present in the surrounding environment^37^. The fact that *HTT* nuclear RNA clusters were disrupted by ammonium acetate underscores the involvement of monovalent cations and ionic interactions in the maintenance of clusters’ integrity and suggests that they might be composed of higher-order RNA assemblies stabilized by secondary and tertiary structures. As ionic bonding also mediates protein-RNA interactions, *HTT* nuclear RNA clusters might be further stabilised by interactions with yet unidentified nuclear protein(s), although the role of PRPF8 cannot be excluded.

### Intermolecular base pairing impacts *HTT* RNA clusters

Building on our hypothesis that *HTT* nuclear RNA clusters could be composed of multiple interacting huntingtin transcripts, we sought to use a treatment that perturbs intermolecular base-pairing, without degrading RNA molecules and doxorubicin emerged as a suitable candidate. Doxorubicin had a striking effect on the number and volume of *HTT* nuclear RNA clusters. Interestingly, it had a biphasic impact on the levels of nuclear and cytoplasmic *FL-HTT* mRNAs which showed an initial increase followed by a drop after 10 hours of treatment. This initial increase might be caused by a greater availability of transcripts after cluster disassembly that would be subjected to nuclear egress. As time lapses doxorubicin might interfere with transcriptional machinery which would result in an overall downregulation of mRNA levels and cluster disappearance. In contrast, nuclear and cytoplasmic *HTT1a* mRNAs showed a sustained, time-sensitive downregulation, consistent with *HTT1a* clusters being reduced more strongly than FL*-HTT* clusters. The mechanism underlying the distinct effects of doxorubicin on FL-*HTT* and *HTT1a* remains unclear. Given that doxorubicin can alter the localization and function of RBPs involved in splicing and polyadenylation^38, 39^, generation of *HTT1a*, which depends on noncanonical processing events, may be particularly vulnerable to such perturbations.

Doxorubicin, however, has pleiotropic effects: its intercalation into DNA double helix causes blockage of transcriptional elongation and inhibition of topoisomerase II impacting chromatin structure^40, 41^. Therefore, future studies are warranted to determine which of these doxorubicin-mediated events lead to disassembly of *HTT* nuclear RNA clusters.

### Nascent transcription nucleates *HTT* RNA clusters

We observed that transcriptional blockage led to a marked decrease in the number of *HTT* nuclear RNA clusters. This was consistent for three different transcription inhibitors, two of which (DRB, NVP-2) specifically hindered the elongation phase. It seems, therefore, that unperturbed *de novo* synthesis of huntingtin mRNA is indispensable for their effective assembly into nuclear clusters. These results also emphasize that *HTT* nuclear RNA clusters are malleable and do not exhibit solid-like behaviour as they can easily undergo internal rearrangements. Interestingly, Pol II transcription processivity is regulated by C-terminal domain phosphorylation, which modifies condensate partitioning behaviour and facilitates redistribution of Pol II from initiation-linked to elongation- and splicing-associated condensates ^32, 42^.

Although our findings suggests that phase separation is not the primary driver of *HTT* nuclear RNA clusters assembly, transcriptional elongation inhibition may partly contribute to dispersion of *HTT* nuclear RNA clusters by disrupting elongation-associated condensates.

Transcriptional inhibition drives relocalisation of numerous RBPs, including TDP-43, from the nucleus to the cytoplasm^23^. If *HTT* RNA clusters engage in prominent interactions with proteins in the nucleoplasm, nuclear efflux of these factors might affect *HTT* nuclear RNA clusters stability. Importantly, in our study, TDP-43 was not identified as an interactor of *HTT* nuclear RNA clusters (Supplementary Fig. 6), however other proteins such as HuR, hnRNPA1, NFX1 also translocate to cytoplasm in response to transcriptional inhibition^23^. Mapping the proteome of *HTT* nuclear RNA clusters will clarify the role of RBPs in maintaining their integrity.

Strikingly, transcriptional inhibition led to intranuclear accumulation of single *FL-HTT* and *HTT*1a mRNAs which increased the number of transcripts amenable to nuclear transport. These single mRNAs likely arise from disintegration of *HTT* nuclear RNA clusters, as transcriptional block would otherwise decrease overall mRNA levels. Importantly, elevated cytoplasmic transcripts could then be translated into toxic proteins, potentially exacerbating cellular dysfunction.

### Conclusions

*HTT* nuclear clusters may have pathogenic or protective implications for cellular health. Conceivably, they could mitigate the production of toxic huntingtin protein isoforms by detaining mutant mRNAs within the nucleus. However, emerging evidence implicates mutant CAG repeat RNA as a pathogenic factor in polyQ disorders. The predominant mechanism of RNA-mediated toxicity involves the formation of stable hairpin structures that sequester pivotal RBPs including MBNL1 leading to deregulated splicing^43, 44^. Indeed, this mechanism underlies disease pathogenesis in myotonic dystrophy (DM1)^43, 45^. Whether *HTT* nuclear RNA clusters interact with MBNL1 remains unresolved as available antibodies failed to yield a detectable signal, leaving this question open for future investigation. In addition to CAG-expanded RNA folding into aberrant secondary structures, recent studies showed that *HTT* transcripts with 40 CAGs undergo constants reengagements^36^. Single stranded regions within these transcripts base-pair with other RNAs promoting formation of larger RNA networks, that may underlie *HTT* nuclear RNA cluster assembly. Such RNA networks might have tremendous consequences for the development of ASO-based therapeutics as these structures could impede ASO hybridization, due to their pairing interactions and 3D geometries. Effective ASO design requires deep insights into RNA conformers and interactors alongside careful consideration of secondary and tertiary RNA topologies.

## Methods

### Ethics statement

All procedures were performed in accordance with the Animals (Scientific Procedures) Act, 1986, and approved by the University College London Ethical Review Process Committee.

### Animal colony breeding and maintenance

YAC128 mice were maintained on a C57BL6/J (Charles River, UK) background. Mice were group-housed depending on gender and genotypes were mixed within cages. All animals were kept in individually ventilated cages containing Aspen Chips 4 Premium bedding (Datesand) with environmental enrichment in the form of chew sticks and a play tunnel (Datesand). All mice had *ad libitum* access to water and chow (Teklad global 18% protein diet, Envigo, The Netherlands). The temperature was automatically regulated at 21°C ± 1°C and animals were kept on a 12 h light / dark cycle. The animal facility was barrier-maintained and quarterly non-sacrificial FELASA (Federation of European Laboratory Animal Science Associations) screens found no evidence of pathogens. Animals were sacrificed at postnatal day 1 and 15 as well as 2 and 12 months of age. Female mice carrying embryos at approximately 14.5 days, were sacrificed at 5 months. The uterine horn was dissected and separated into individual embryos and embryos were released from the embryonic sac. Thereafter, the head of the embryo was cut to separate it from the body; a tail cut was taken into a separate Eppendorf for genotyping. Two templar incisions were made by inserting the tips of the scissors as to not cut through the brain; then, watchmakers were used to gently lift the skull and separate it from the brain. Once the brain was exposed, it was lifted from below to remove it from the head. This process was repeated for all YAC128 animals sacrificed, however from P15 onwards the brains were separated into hemispheres which were then embedded individually in Optimal cutting temperature (OCT) compound (Cell Path, KMA-0100-00A).

### DNA extractions, genotyping and repeat sizing

Genomic DNA from embryonic day 14.5 animals and postnatal day 1 pups was extracted from the tail cut. Genomic DNA from postnatal day 15 pups and 2,12-month-old mice was obtained from ear notches and genotyping was performed as previously described^46^. The polyQ repeat of 125 glutamines in YAC128 mice is encoded by (CAG)_23_(CAA)_3_CAGCAA(CAG)_80_(CAA)_3_CAGCAA(CAG)_10_CAACAG which is stable on germline transmission.^47^

### Primary neuronal cultures

For the preparation of primary hippocampal cultured cells derived from neonatal YAC128 mice and their wild-type siblings, postnatal animals (day P0-P2) were sacrificed by decapitation. Subsequently, hippocampi were isolated, deprived of meninges and collected into 2 mL Eppendorf containing dissection medium (HBSS (Gibco, 14175-053) supplemented with 1x sodium pyruvate (Gibco, 11360-070), 20% glucose (Sigma,G8270-1KG),1M HEPES pH=7.3 (Gibco,15630-080), thereafter hippocampi were washed twice with dissection medium and 1 mL of papain solution containing 60 µL of DNase was added (per two hippocampi; Papain dissociation system kit, Worthington Chemical Corporation, LK003150). After 15 min incubation at 37°C, tissue was washed once with warm dissection medium and twice with warm plating medium (DMEM (Gibco, 41966-029) supplemented with 10% FBS (Gibco, 16140-071), 20% glucose, 1x sodium pyruvate,1x glutamax (Gibco, 35050-061) and penicillin/streptomycin (Gibco, 15070-063)) and then subjected to mechanical tituration first with 1 mL pipette and then with 100 µL pipette. In the next step, tissue was centrifuged at 100 rcf for 5 min at RT, the supernatant was discarded, and cells resuspended in 670 µL EBBS with 75 µL albumin-ovomucoid inhibitor and 35 µL DNase (per animal, Papain dissociation system kit). The mixture was layered atop 1.25 mL of albumin-ovomucoid inhibitor and centrifuged at 1880 rcf for 6 min at RT. The cell pellet was resuspended in 500 µL of plating medium, and equal volumes of the resuspended cells and Trypan Blue solution were mixed. Cell viability and number were then determined using the Tecan cell counting method.Thereafter, cells were diluted into plating medium and plated onto poly-D-lysine-coated (R&D systems, 3439-100-01) glass coverslips (VWR, 631-0149), placed into 24 well plate (Falcon, 353947). After 2 h, 1 mL of maintenance medium (Neurobasal medium (Gibco, 21103-049) supplemented with B27 (Gibco, A35828-01), 1x glutamax and penicillin/streptomycin) was added to each well of a 24-well plate and placed in a humidified incubator. Cells were plated at a density of 150, 000/ coverslip and cultured for 18-10 days *in vitro* (DIV) unless stated otherwise.

### RNAscope analysis and quantification

Whole brains (embryonic day 14.5; postnatal day 1 and 15) or brain hemispheres (2,12 months) from YAC128 mice were harvested, embedded in optimal cutting temperature compound (OCT, CellPath Ltd) and stored at −80°C until further use. Brain blocks were equilibrated to about −30°C in a cryostat (Bright Instruments), cut into 20 µm thick sagittal sections and mounted onto SuperfrostPlus slides (VWR, 25 x 75 x 1.0 mm). The brain sections, or cells cultured on coverslips, were fixed in 4% paraformaldehyde (Pioneer Research Chemical Ltd,) for 20-30 min at 4°C followed by dehydration in an increasing concentration of ethanol, starting at 50%, moving to 70% and finishing with 100%, each time for 5 min (for brain sections) or 1 min (for cultured cells) at RT. Next, a hydrophobic barrier was created around each section with a hydrophobic barrier pen (ImmEdge Pen, Vector; H-4000) and allowed to air-dry for about 1 min. About five drops of 3% hydrogen peroxide solution (H_2_O_2_) was added onto the section and incubated for 10 min at RT, followed by three washes with PBS. Protease IV (for brain sections) or protease III (for cultured cells, diluted 1:20) was then added to the samples and incubated for 30 min (for brain sections of animals at 2 and 12-months of age), 15 min (for brain sections of animals at embryonic day 14.5; postnatal days 1 and 15) or 10 min (for cultured cells) at RT. The protease was removed, and the slides submerged in PBS three times for 5 min each.

The hybridization and amplification steps were performed in accordance with the RNAscope protocol (Advanced Cell Diagnostics, Inc.). Briefly, probes were hybridized for 2 h at 40°C in a HybEZ oven (Advanced Cell Diagnostics, Inc., 220 VAC, 310013). Following each amplification step, the slides were washed twice with PBS for 5 min each at room temperature. Each probe was developed in three steps: the HRP (horse radish peroxidase) step, the TSA (tyramide signal amplification) fluorophore step and the HRP blocker step, using the RNAscope Multiplex Fluorescent v2 Assay. Each step was followed by two washes in PBS for 5 min at RT. A different TSA fluorophore (PerkinElmer) was assigned to each probe channel. The following fluorophores were used at 1:1500 dilution: fluorescein, Cy3 and Cy5. Subsequently, the slides were counterstained with DAPI for about 5 min at RT. Tissue section slides were mounted in VECTASHIELD Vibrance Antifade Mounting Media (Vector Laboratories, H-1000-10), a 24 x 60 mm glass coverslip (VWR International, ECN 631-1575) was placed over the tissue section, and the slides were air dried before being imaged. Cells on the coverslips were mounted in EverBrite ^TM^ hardset mounting medium (Biotium, 23003) and air-dried before imaging. Images were acquired with an inverted (Zeiss LSM980) confocal integrated microscope system with a 63x objective (HC PL APO 63x/1.40 Oil).

### Immunofluorescence

Immunofluorescence was conducted in conjunction with the RNAscope protocol. For combined RNAscope-IF experiments, primary antibodies were diluted 1:10 in Co-Detection Diluent (Advanced Cell Diagnostics, 323160) and were applied after the hydrogen peroxidase step and incubated overnight at 4°C. Primary antibodies included: NeuN (Abcam, ab177487), GFAP (Invitrogen, PA1-10004), PrPF8 (Abcam, ab79237), coilin (Abcam, ab11822), SFPQ (Abcam, ab38148), H3K9ac (Upstate, cell signalling solutions06-942), p84 (GeneTex, GTX118740), TDP-43 (Abcam, ab104223), hnRNP K (Abcam, ab52600), hnRNP F (Thermofisher Scientific, CF500804). The following day cells were washed twice with PBS and subjected to the regular RNAscope procedure described above. Before DAPI staining, secondary antibodies were applied at RT for 2 hours.

Secondary antibodies were diluted in Co-Detection Diluent, used at 1:200 and included anti-mouse (Alexa555, Invitrogen, A32773), ani-rabbit (Alexa555, Invitrogen, A32794), anti-goat (Alexa488, Invitrogen, A11055).

### Hippocampal cell treatments

5% or 8% 1,6-Hexanediol (Sigma, 240117-50G) and 8% 2,5-Hexanediol (Merck, 240117-50MG) or an equivalent dilution of water as control were added to hippocampal cells in culture for 5 min. Ammonium acetate (Sigma, A1542-250G) was added to the cultured media and cells were incubated in this medium for 10 min. Cells were treated with transcriptional inhibitors: ActinomycinD (Sigma, A1410-2MG), NVP2 (Tocris, 6535) and DRB (Millipore, 287891-50mg) for 3 h. Splicing blocker, PladB (Cayman Chemical company, 364-9897) was added to hippocampal cells in culture for 4 h. Cells were incubated with doxorubicin (Cell signalling Technology, 5927S) for either 5 or 10 h. Unless otherwise stated DMSO (Sigma, D2438-10ML) served as a control.

### Analysis of RNA clusters/ transcripts

We used a size-based threshold to identify RNA clusters/puncta. Briefly, YAC128 sections were imaged using a confocal microscope, and 1 μm Z-stacks were acquired. For YAC128 hippocampal cells, Z-stacks at 0.5 μm were acquired. Images were converted to 8-bit and a DAPI signal was used to create a threshold-based mask for cell nuclei to separate nuclear and cytoplasmic signals for different huntingtin transcripts. To create a Z-stack of molecules outside of the nucleus an 8-bit Z-stack containing images of one of the three *HTT* transcripts was subtracted from the binary DAPI mask Z-stack. To generate a Z-stack of molecules inside of the nucleus an 8-bit Z-stack containing images of one of the three *HTT* transcripts was subtracted from the previously created out-of-the-nucleus Z-stack. RNA clusters and nuclear puncta as well as cytoplasmic mRNA puncta were identified using FIJI 3D Objects Counter plugin^48^. An intensity threshold in this plug-in was set to segment objects from the background, with intensity threshold varying between different treatments but kept constant for control and treatment within the same experimental group. Thereafter, the size filter was set to the minimum of about 10 voxels to filter out noise, and the maximum was set automatically by the programme to include large nuclear structures. This plug-in faithfully identified clusters/single mRNAs as depicted in Supplementary Fig. 2a. The method was applied to determine the number, volume and surface area of the clusters/ single mRNAs. The number as well as the total volume taken up by the clusters per cell were chosen as the parameters to quantify the extent of cluster formation.

### Statistical analysis

The results are expressed as the mean□±□SEM. To test the data for normal distribution D’Agostino-Pearson test was applied to the developmental YAC128 data and Shapiro-Wilk test was applied to all data collected from the hippocampal YAC128 cell cultures. Whenever normality requirements were not met, a non-parametric test was used. For experiments determining developmental trajectory of clusters appearance either one-way ANOVA followed by a post-hoc Bonferroni’s test (parametric) or Kruskal-Wallis with Dunn’s correction for multiple comparisons (non-parametric) were used. For experiments on hippocampal cells comparing the effect of different pharmacological agents on the clusters/transcripts number/volume, statistical significance was analysed using either unpaired, two-tailed *t*-test or unpaired, two-tailed Mann-Whitney tests. Statistical analyses were performed using GraphPad Prism 10 (GraphPad Software Inc.). The statistical tests used and sample size for each analysis are shown in the Figure legends.

## Data availability

All data generated or analysed during this study are included in this publication and its supplementary information. Raw data will be shared by the corresponding author upon request.

## Contributions

S.F and G.B conceived the study. S. F. drafted the manuscript. S.F, I.N and I.M.M. completed the data collection and S.F carried out the analysis. G.B provided funding. All authors critically reviewed the manuscript and agreed to submit it for publication.

## Corresponding authors

Correspondence to Sandra Fienko.

