## Supplementary material for "Nuclear RNA Clusters Are Dynamic Structural Entities in Huntington’s Disease": Fienko et al._2025_Supplement.pdf

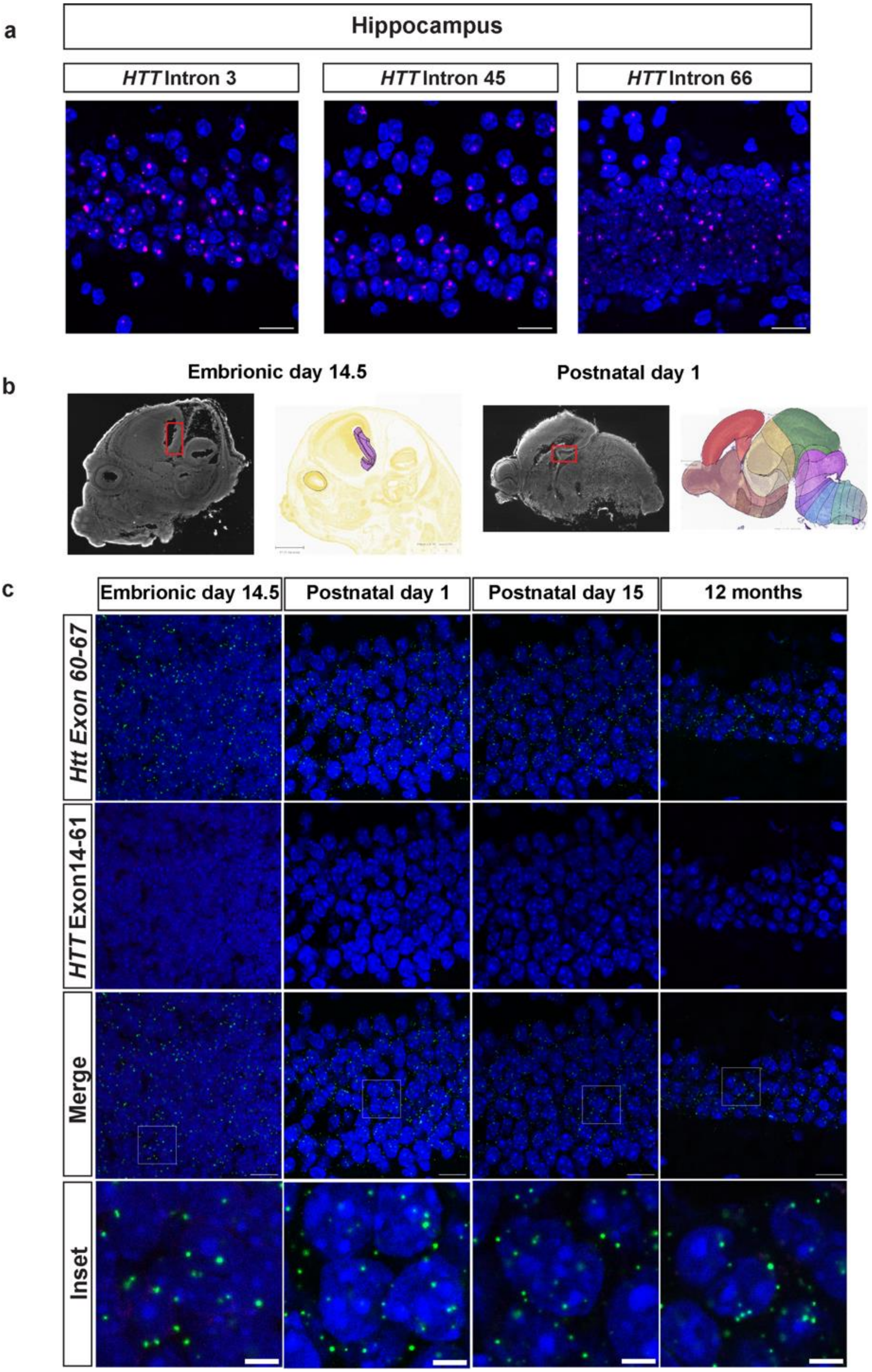
a
Hippocampus
HTT Intron 3
HTT Intron 45
HTT Intron 66
b
Embrionic day 14.5
Postnatal day 1
c
Embrionic day 14.5
Postnatal day 1
Postnatal day 15
12 months
Htt Exon 60-67
HTT Exon14-61
Merge
Inset

**Supplementary Figure 1 Different intronic *HTT* probes revealed different levels of nuclearly enriched *HTT* pre-mRNA in the (developing) hippocampus.**

**(a)** Hippocampal sections from YAC128 mice at 2 months of age were hybridised with different intronic huntingtin probes: *HTT*inton 3, *HTT*inton 45 and *HTT*inton 66. Images revealed that *HTT*inton 3 and *HTT*inton 45 probes detected similar levels of unprocessed huntingtin message, whereas *HTT*inton 66 probe underestimated the levels of huntingtin pre-mRNA. Nuclei were stained with DAPI (blue). Scale bar is 20 µm in the main image

**(b)** Representative images showing the region of interest (red square box) imaged under confocal microscopy corresponding to the precursor of hippocampal formation, medium pallium and the matching region in the Allen Brain Atlas at earlies developmental stages: embryonic day 14.5 and post-natal day 1.

**(c)** Sagittal sections from wild-type brains at embryonic day E14.5, postnatal days P0 and P15 as well as 12 months of age were hybridised with probes to human and mouse full-length huntingtin transcripts. Sections show no staining for FL-*HTT* (human, *HTT* Exon 14-61), whereas FL-*Htt* (mouse, *Htt* Exon 60-67) is detected in the nucleus and in the cytoplasm. Nuclei were stained with DAPI (blue). Wild type (*n* = 3). Scale bar = 20 µm in the main image and 5 µm in the cropped magnified image.

a

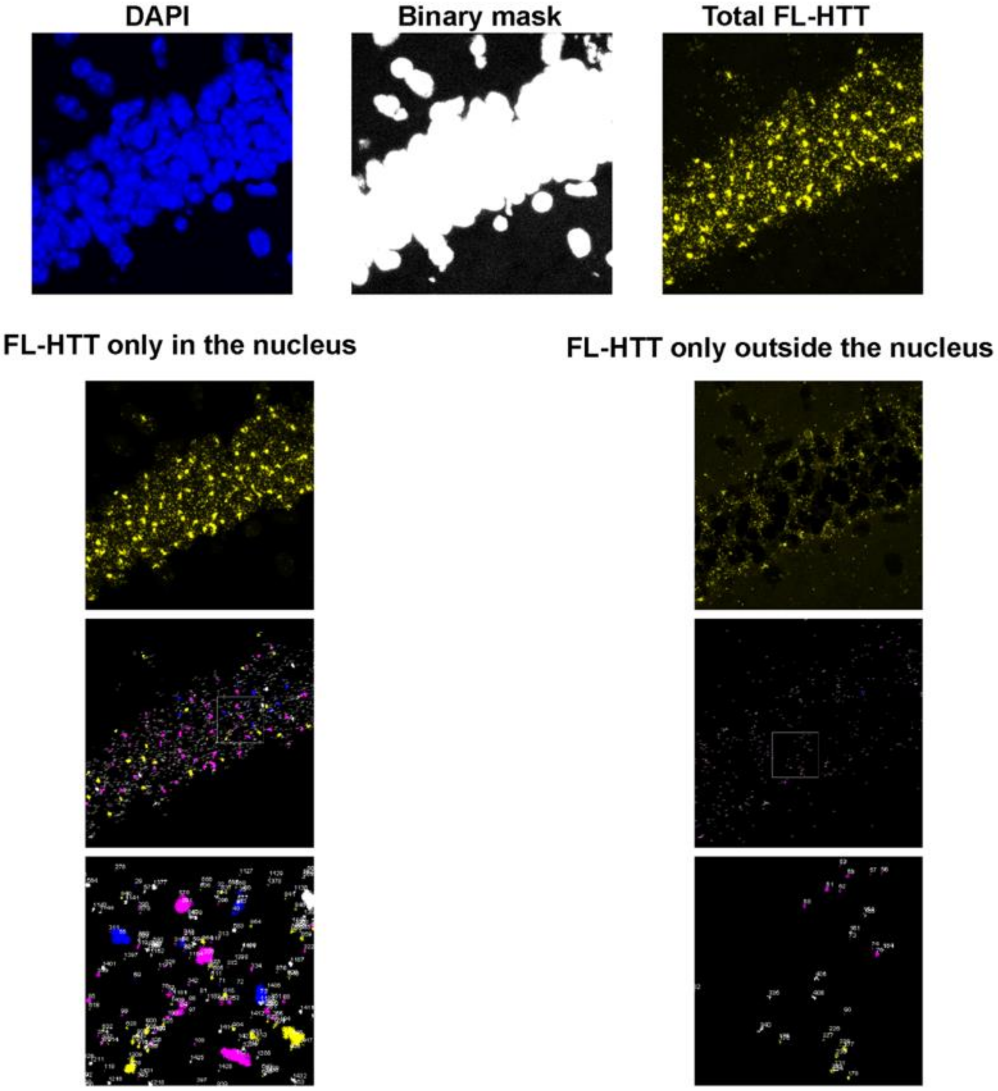
DAPI
Binary mask
Total FL-HTT
FL-HTT only in the nucleus
FL-HTT only outside the nucleus

b

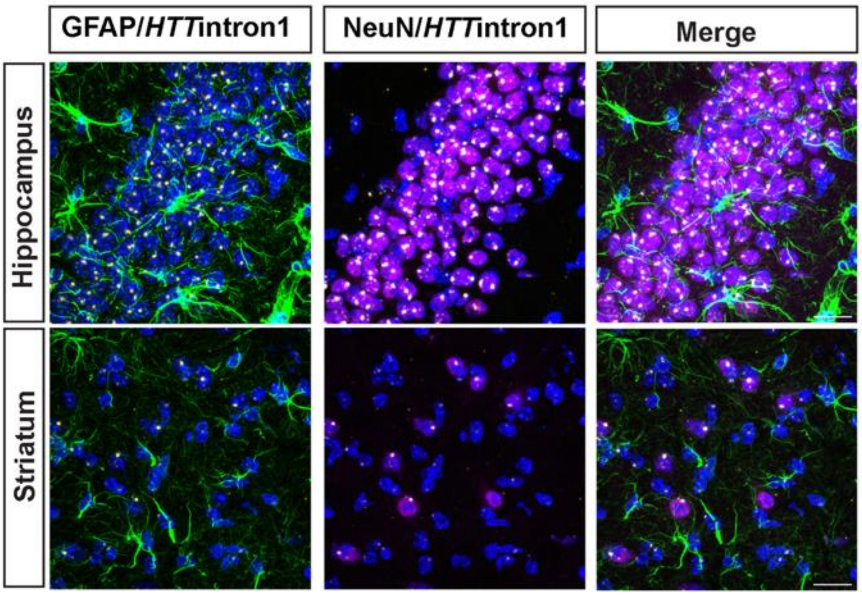
GFAP/*HTT*intron1
NeuN/*HTT*intron1
Merge
Hippocampus
Striatum

**Supplementary Figure 2 *HTT* nuclear RNA clusters were localised to neurons and were not detected in the brains of wild-type mice.**

**(a)** To identify RNA clusters/foci size-based threshold was used. DAPI signal was used to establish a threshold-based mask for cell nuclei to isolate nuclear and cytoplasmic signals for different huntingtin transcripts. To obtain a Z-stack of cytoplasmic molecules, the Z-stack of *HTT* transcript images was subtracted from the DAPI mask Z-stack. A nuclear Z-stack was then generated by subtracting the cytoplasmic Z-stack from the original *HTT* transcript Z-stack. RNA clusters and nuclear puncta as well as cytoplasmic mRNAs were analysed with FIJI 3D Objects Counter plugin. Intensity threshold was set to isolate objects from the background. As illustrated the plug-in faithfully identified clusters/single mRNAs.

**(b)** Hippocampal and striatal sections from YAC128 mice at 2 months of age were stained with the NeuN (neuronal marker) antibody and a GFAP antibody to visualize astrocytes. Thereafter, sections were hybridized with RNAscope probes against intron 1 human *HTT* sequences. *HTT* nuclear RNA clusters were prevalent in neuronal cells. Scale bar is 20 μm.

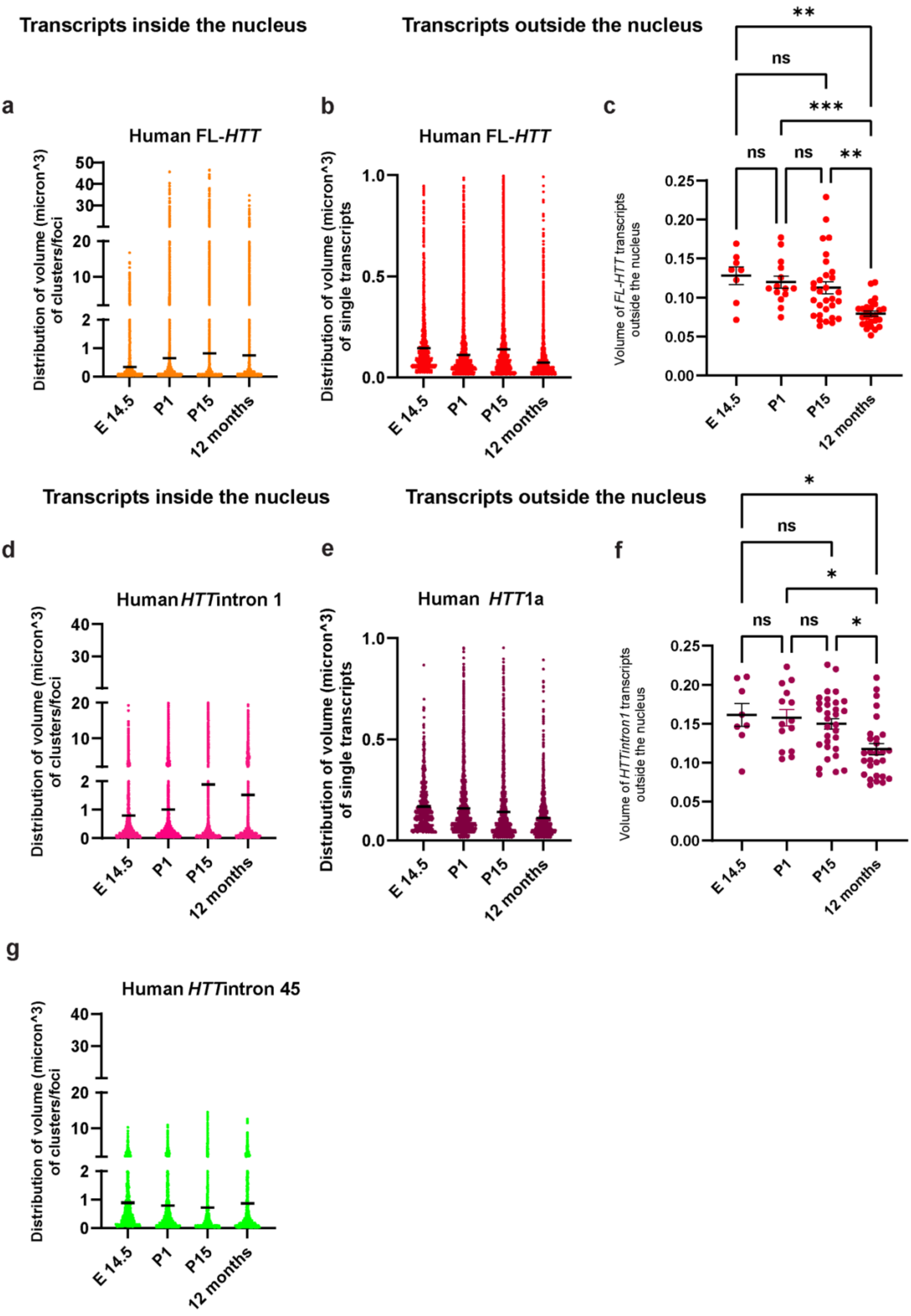

Transcripts inside the nucleus
Transcripts outside the nucleus
a
Human FL-*HTT*
Distribution of volume (micron^3) of clusters/foci
50
40
30
20
10
2
1
0
E 14.5
P1
P15
12 months
b
Human FL-*HTT*
Distribution of volume (micron^3) of single transcripts
1.0
0.5
0.0
E 14.5
P1
P15
12 months
c
**
ns
***
ns
ns
**
Volume of *FL-HTT* transcripts outside the nucleus
0.25
0.20
0.15
0.10
0.05
0.00
E 14.5
P1
P15
12 months
Transcripts inside the nucleus
Transcripts outside the nucleus
d
Human *HTT* intron 1
Distribution of volume (micron^3) of clusters/foci
40
30
20
10
2
1
0
E 14.5
P1
P15
12 months
e
Human *HTT*1a
Distribution of volume (micron^3) of single transcripts
1.0
0.5
0.0
E 14.5
P1
P15
12 months
f
*
ns
*
ns
ns
*
Volume of *HTTintron1* transcripts outside the nucleus
0.25
0.20
0.15
0.10
0.05
0.00
E 14.5
P1
P15
12 months
g
Human *HTT* intron 45
Distribution of volume (micron^3) of clusters/foci
40
30
20
10
2
1
0
E 14.5
P1
P15
12 months

**Supplementary Figure 3 Distribution of the volume of *HTT* nuclear RNA clusters and single mRNA transcripts inside and outside the nucleus.**

Distribution of the total volume of all objects detected by 3D-volume analysis plug-in in ImageJ. Objects correspond to either nuclear clusters or mRNA transcripts, and these objects were quantified for **(a)** FL-*HTT* probe inside the nucleus, whereas **(b)** depicts single mRNA transcripts outside the nucleus for FL-*HTT* probe. **(c)** Averaged volume of FL-*HTT* transcripts outside the nucleus across YAC128 development (E14.5=0.13±0.01, P1=0.12±0.008, P15=0.12±0.008, 12-months=0.08±0.003). **(d)** Objects corresponding to either nuclear clusters or single intranuclear mRNA transcripts are indicated for *HTT*intron 1 probe. **(e)** The volume of the transcripts in the cytoplasmic compartment was also plotted for *HTT*intron 1 probe. **(f)** Quantification of averaged volume of extranuclear *HTT*1a mRNAs at various developmental stages (E14.5=0.16±0.01, P1=0.16±0.01, P15=0.15±0.7, 12-months=0.12±0.007). YAC128 (n = 3). **(g)** Distribution of the total volume of all objects present in the nucleus for *HTT*inton 45 probe.

D'Agostino-Pearson test was applied to test data for normal distribution and Kruskal-Wallis test was used to verify significance of data. Error bars = mean ± SEM, *$P \leq 0.033$, **$P \leq 0.002$, ***$P \leq 0.0002$, ****$P \leq 0.0001$.

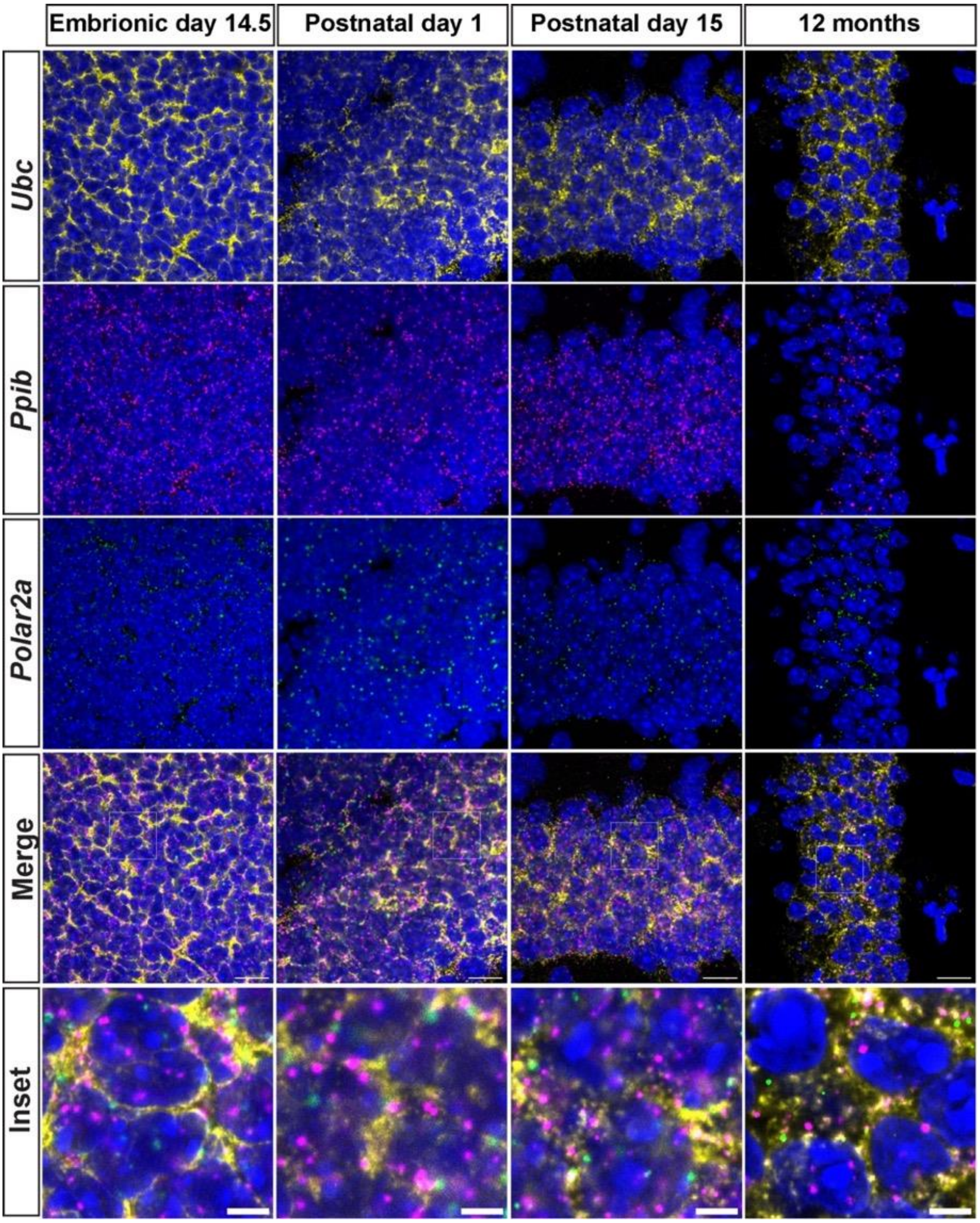


**Supplementary Figure 4 Detection of mRNA transcripts for three differentially expressed housekeeping genes in the brains of YAC128 mice at different developmental stages.**

Sagittal sections from YAC128 animals at embryonic day E14.5, postnatal days P0 and P15 as well as 12 months of age were stained with a set of probes recognising housekeeping transcripts. *Ubc* (yellow) is a highly expressed mRNA, *Ppib* (magenta*)* is expressed at high to medium levels whereas *Polar2a* (green) is a low abundant mRNA. Note that most of the transcripts reside outside of the nucleus. Nuclei were stained with DAPI (blue). YAC128 ($n$ = 3), Scale bar = 20 µm in the main image and 5 µm in the cropped magnified image.

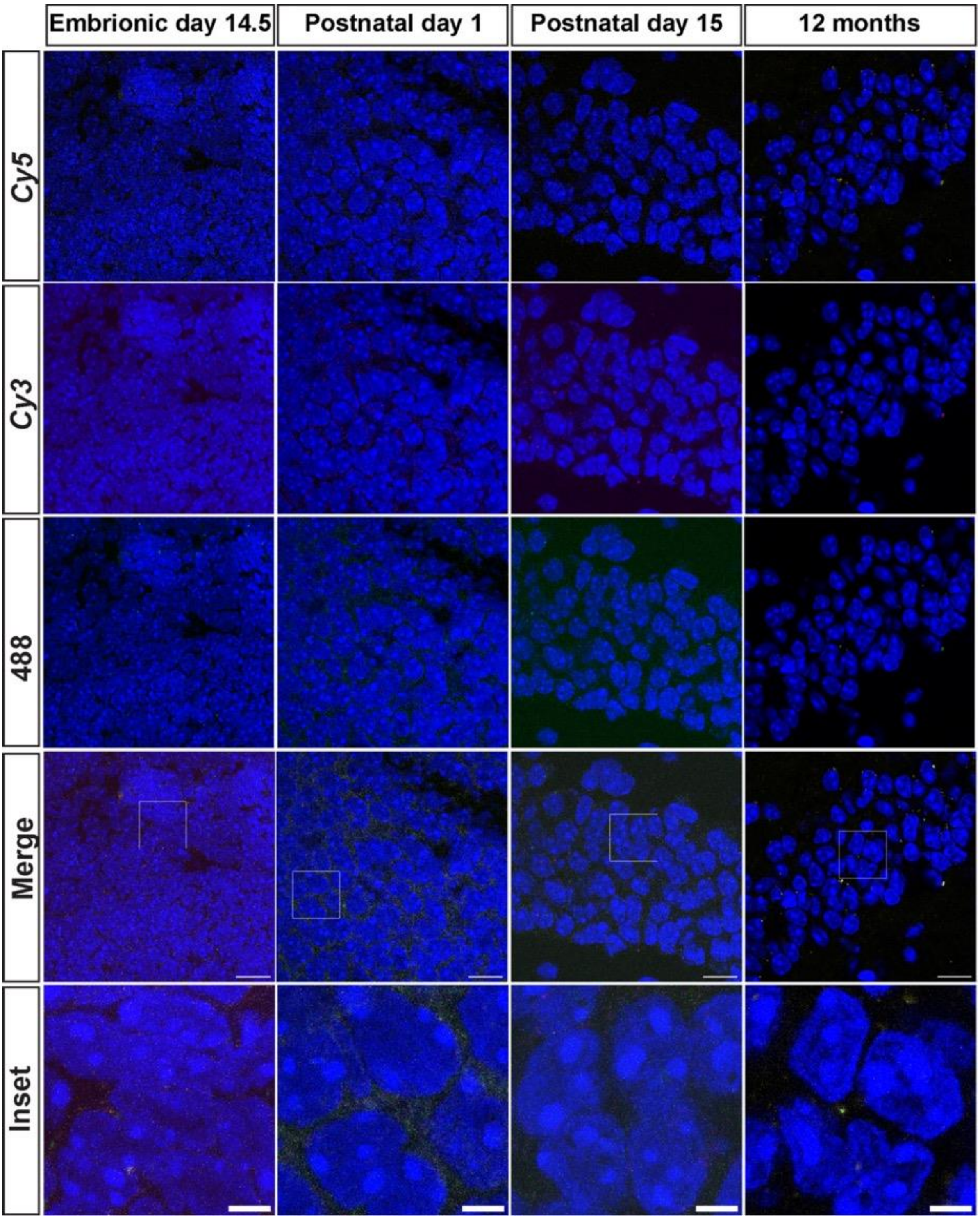


**Supplementary Figure 5 Sections from YAC128 mice at different developmental stages showed no staining when hybridised with probes against a bacterial gene that served as a negative control.**

Sagittal sections from YAC128 animals at embryonic day E14.5, postnatal days P0 and P15 as well as 12 months of age were stained with negative control probes targeting the DapB gene from the Bacillus subtilis strain SMY labelled with each of the

Cy5, Cy3 and 488 fluorophores. No background staining was observed in any of the channels. Nuclei were stained with DAPI (blue). YAC128 (*n* = 3), Scale bar = 20 μm in the main image and 5 μm in the cropped magnified image.

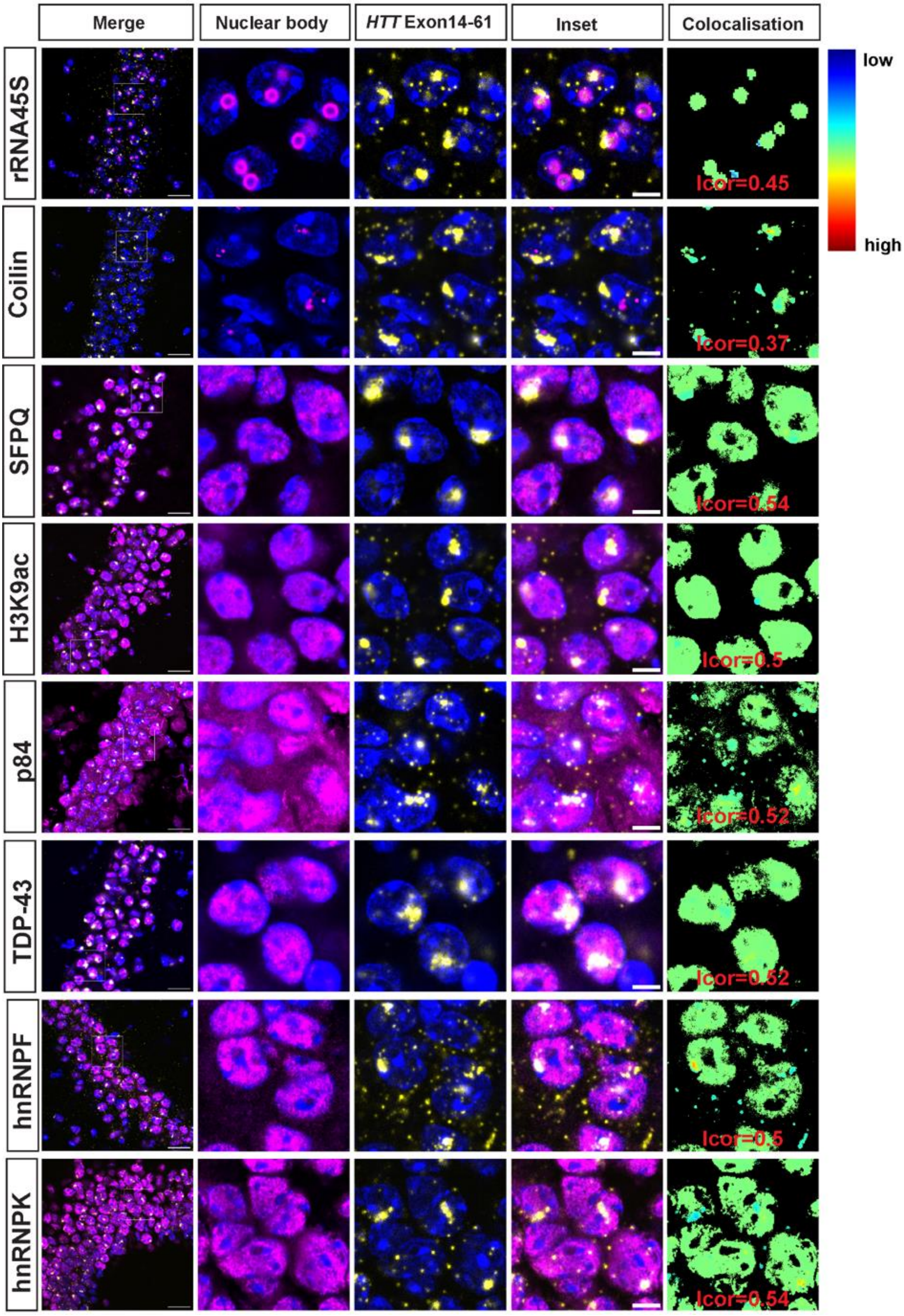
Merge
Nuclear body
HTT Exon14-61
Inset
Colocalisation
low
high
rRNA45S
Icor=0.45
Coilin
Icor=0.37
SFPQ
Icor=0.54
H3K9ac
Icor=0.5
p84
Icor=0.52
TDP-43
Icor=0.52
hnRNPF
Icor=0.5
hnRNPK
Icor=0.54

**Supplementary Figure 6 *HTT* nuclear RNA clusters do not colocalise with the markers of well-identified nuclear bodies.**

Representative immunofluorescence micrographs depicting that *HTT* nuclear RNA clusters (revealed in yellow by *HTT*exon14-61 probe) do not associate with the marker for nucleolus (rRNA45S), Cajal bodies (coilin), paraspeckles (SFPQ), open state chromatin (H3K9ac), RNA nuclear export (p84) or various RNA-binding proteins (TDP-43, hnRNPF, hnRNPK). Icor represents the 'Index of correlation' and indicates the extent of colocalization. Analysis was performed using ImageJ plug-in Colocalisation Colormap.

Nuclei were stained with DAPI (blue). YAC128 (*n* = 3), Scale bar = 20 μm in the main image and 5 μm in the cropped magnified image.

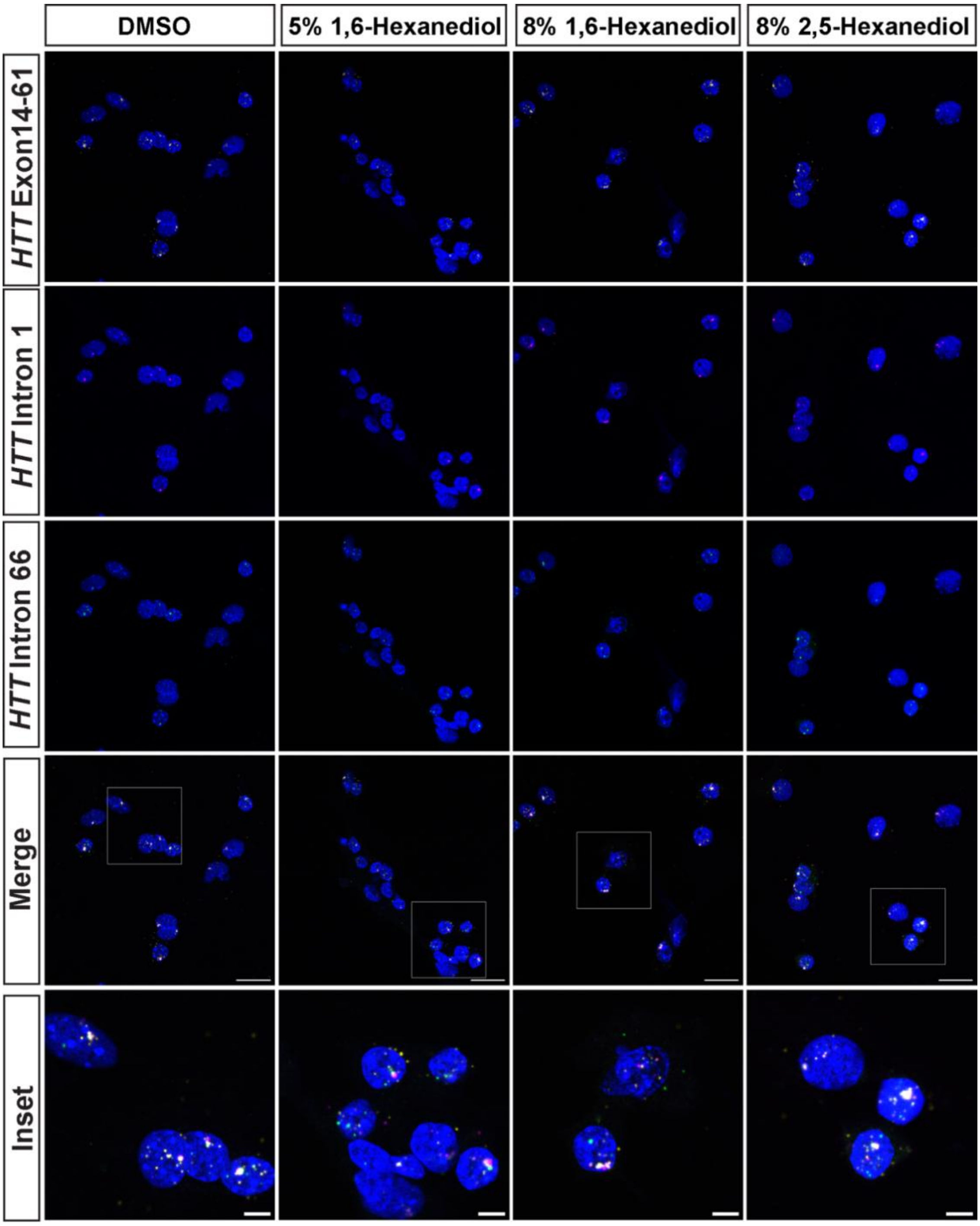


**Supplementary Figure 7 *HTT* nuclear RNA clusters are not visibly disrupted by agents that dissolve liquid-liquid phase separated condensates**.

Schematic representation of 18-20 DIV hippocampal cultured cells treated with either 5% or 8% 1,6-Hexanediol and less potent 8% 2,5-Hexanediol. The number of *HTT* nuclear RNA clusters was not visibly altered. FL-*HTT* (yellow), *HTT*intron 1 (magenta), *HTT*intron 66 (green), nuclei were stained with DAPI (blue). Number of individual cultures n = 3, number of transgenic mouse pups per culture imaged n= 1. Scale bar is 20 μm in the main image and 5 μm is the cropped magnified image.

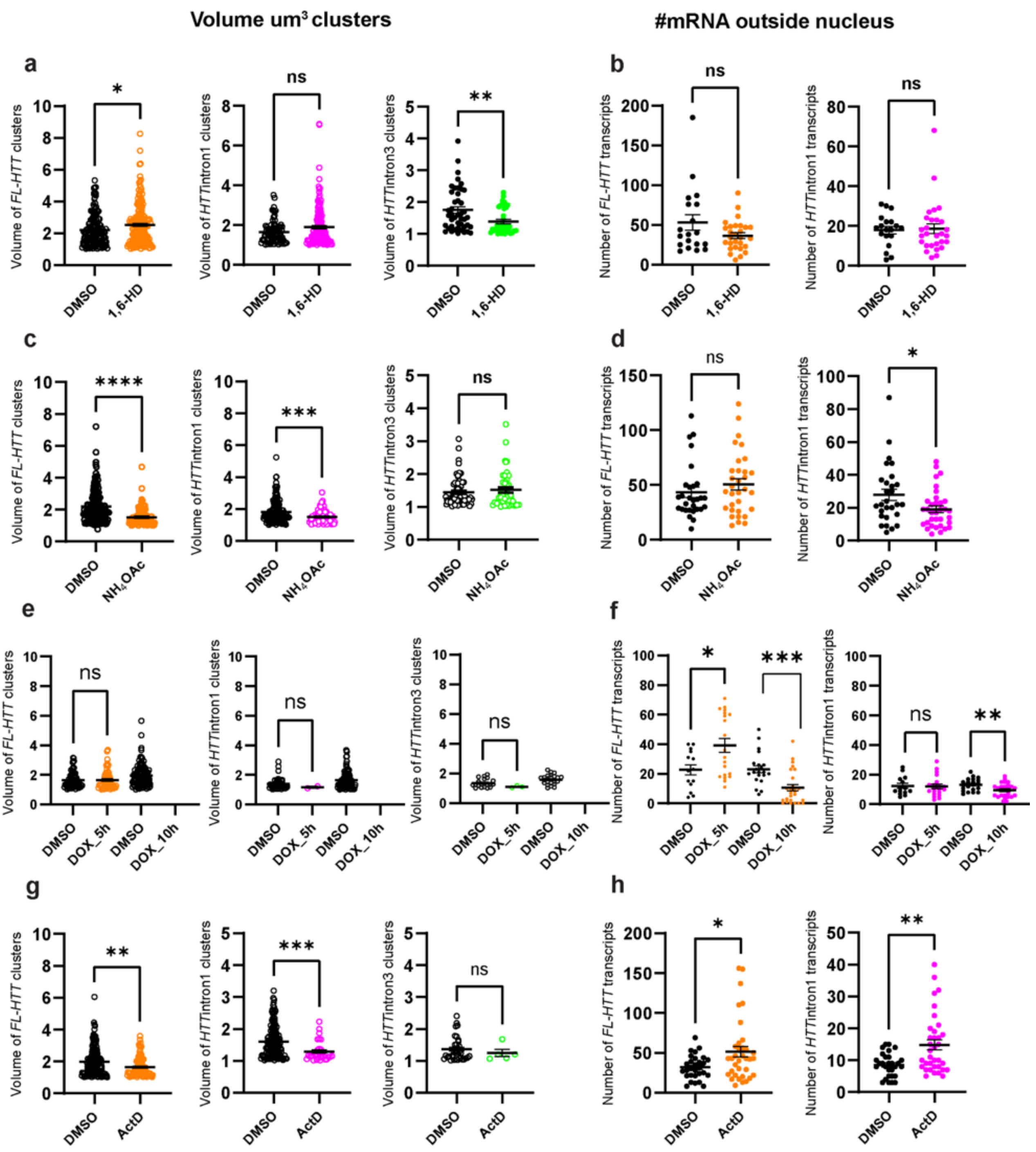

Volume um³ clusters
#mRNA outside nucleus
a
b
c
d
e
f
g
h
Volume of FL-HTT clusters
Volume of HTTintron1 clusters
Volume of HTTintron3 clusters
Number of FL-HTT transcripts
Number of HTTintron1 transcripts
DMSO
1,6-HD
NH4OAc
DOX_5h
DOX_10h
ActD
ns
*
**
***
****

**Supplementary Figure 8 Quantification of the volume of *HTT* nuclear RNA clusters and the number of RNA huntingtin transcripts upon various pharmacological treatments.**

Analysis of the volume of *HTT* nuclear RNA clusters was performed upon treatments with **(a)** 1,6-Hexanediol (1,6-HD) (FL-*HTT*: 1,6-HD=2.5±0.09, DMSO=2.2±0.08; *HTT*intron 1: 1,6-HD=1.9±0.09, DMSO=1.6±0.06; *HTT*intron 3: 1,6-HD=1.4±0.07, DMSO=1.7±0.1), **(c)** 100 mM ammonium acetate (FL-*HTT*: $NH_4OAc$=1.5±0.05, DMSO=2.2±0.07*;* *HTT*intron 1: $NH_4OAc$=1.5±0.03, DMSO=1.8±0.06; *HTT*intron 3: $NH_4OAc$=1.5±0.08, DMSO=1.4±0.05)**,** **(e)** 2 μM doxorubicin (5-hours: FL-*HTT*: Dox=1.6±0.08, DMSO=1.6±0.06; *HTT*intron 1: Dox=1.2±0.06, DMSO=1.4±0.06; *HTT*intron 3: Dox=1.1±0.03, DMSO=1.3±0.06; 10-hours: FL-*HTT*: Dox=0.0±0.0, DMSO=1.9±0.07; *HTT*intron 1: Dox=0.0±0.0, DMSO=1.6±0.06; *HTT*intron 3: Dox=0.4±0.0, DMSO=1.6±0.06), **(g)** 1 μM Actinomycin D (FL-*HTT*: ActD=1.6±0.06, DMSO=2±0.07, *HTT*intron 1: ActD=1.3±0.06, DMSO=1.6±0.04, *HTT*intron 3: ActD=1.2±0.1, DMSO=1.4±0.06).

The number of the huntingtin transcripts outside of the nucleus was established for the following agents **(b)** 1,6-HD (FL-*HTT*: 1,6-HD=37±4, DMSO=53±9; *HTT*1a: 1,6-HD=19±2, DMSO=18±2)., **(d)** 100 mM ammonium acetate (FL-*HTT*: $NH_4OAc$=51±5, DMSO=43±5; *HTT1a*: $NH_4OAc$=19±2; DMSO=28±3), **(f)** 2 μM doxorubicin (5-hours: FL-*HTT*: Dox=39±4, DMSO=23±3; *HTT*1a: Dox=12±1, DMSO=12±2; 10-hours: FL-*HTT*: Dox=11±2, DMSO=23±2; *HTT*1a: Dox=9±1, DMSO=13±1), **(h)** 1 μM Actinomycin D (FL-*HTT*: ActD=51±6, DMSO=32±3, *HTT*1a: ActD=15±2, DMSO=9±0.7). Number of individual cultures n = 3, number of transgenic mouse pups per culture imaged n= 2. Scale bar is 20 μm in the main image and 5 μm is the cropped magnified image. Shapiro-Wilk test was applied to test data for normal distribution and either Mann-Whitney test (all treatments: volume of the clusters for all three probes; number of the transcripts outside the nucleus for FL-*HTT* and *HTT*1a for 1,6-HD, $NH_4OAc$, ActD and for FL-*HTT* for doxorubicin treatment) or unpaired two-tailed *t*-test (number of the transcripts outside the nucleus for *HTT*1a for doxorubicin) was subsequently used to verify the significance of data. Error bars = mean ± SEM, **P* ≤ 0.033, ***P* ≤ 0.002, ****P* ≤ 0.0002, *****P* ≤ 0.0001.

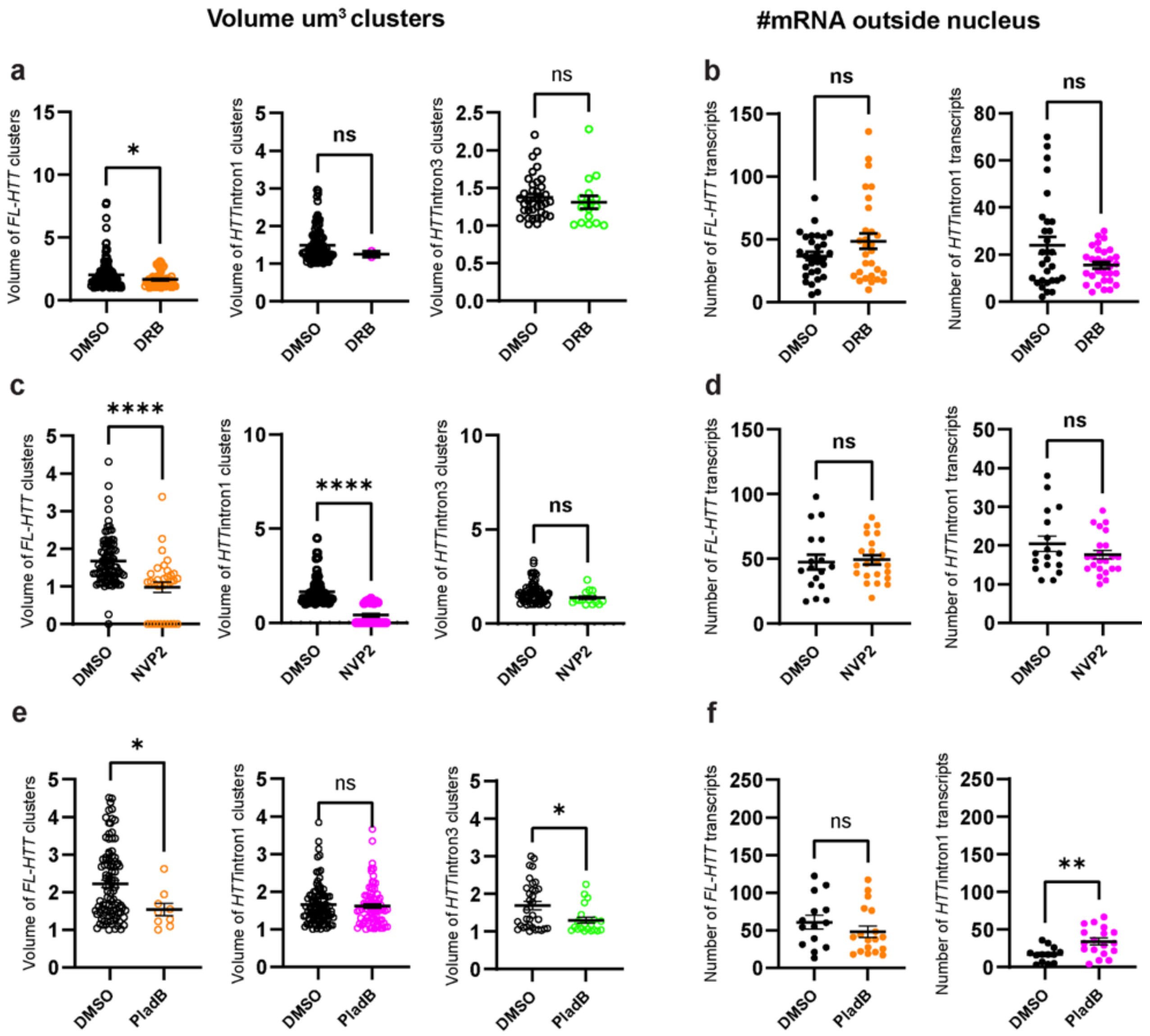


**Supplementary Figure 9 Quantification of the volume of *HTT* nuclear RNA clusters and the number of RNA huntingtin transcripts upon transcriptional elongation inhibitors and PladB treatments.**

Analysis of the volume of *HTT* nuclear RNA clusters upon application of **(a)** 100 μM DRB (FL-*HTT*: DRB=1.6±0.07, DMSO=2±0.1; *HTT*intron 1: DRB=1.3±0.07, DMSO=1.5±0.04; *HTT*intron 3: DRB=1.3±0.08, DMSO=1.4±0.05). **(c)** 250 nM NVP-2 FL-*HTT*: NVP-2=1±0.1, DMSO=1.7±0.07; *HTT*intron 1: NVP-2=0.4±0.1, DMSO=1.7±0.08; *HTT*intron 3: NVP-2=1.4±0.09, DMSO=1.6±0.07) and **(e)** the splicing inhibitor PladB (FL-*HTT*: PladB=1.5±0.1, DMSO=2.2±0.09; *HTT*intron 1: PladB=1.6±0.06, DMSO=1.6±0.05; *HTT*intron 3: PladB=1.3±0.08, DMSO=1.6±0.1) The number of the huntingtin transcripts outside of the nucleus was quantified for **(b)** 100 μM DRB (FL-*HTT*: DRB=49±6, DMSO=37±3; *HTT1a*: DRB=15±1, DMSO=24±4) and **(d)** 250 nM NVP-2 (FL-*HTT*: NVP-2=49±3 DMSO=48±6, *HTT1a*: NVP-2=18±1, DMSO=20±2) and **(f)** PladB (FL-*HTT*: PladB=48±7 DMSO=61±9; *HTT1a*: PladB=34±4, DMSO=16±3). Number of individual cultures n = 3, number of transgenic mouse pups per culture imaged n= 2. Scale bar is 20 µm in the main image and 5 µm is the cropped magnified image. Shapiro-Wilk test was applied to test data for normal distribution and either Mann-Whitney test (all treatments: volume of the clusters for all three probes; number of the transcripts outside the nucleus for *HTT*intron 1 for DRB and NVP-2 and FL-*HTT* for DRB and PladB) or unpaired two-tailed *t*-test ( number of the transcripts outside the nucleus for *HTT*intron 1 for PladB and FL-*HTT* for NVP-2) was subsequently used to verify the significance of data. Error bars = mean ± SEM, **P* ≤ 0.033, ***P* ≤ 0.002, ****P* ≤ 0.0002, *****P* ≤ 0.0001.
